# Using runs of homozygosity and machine learning to disentangle sources of inbreeding and infer self-fertilization rates

**DOI:** 10.1101/2024.02.20.581206

**Authors:** Leo Zeitler, Kimberly J. Gilbert

**Affiliations:** Department of Biology, University of Fribourg, Chemin du Musée 10, 1700 Fribourg, Switzerland

**Keywords:** runs of homozygosity, inbreeding, self-fertilization, outcrossing rate, demographic history, mating system, random forest

## Abstract

Runs of homozygosity (ROHs) are indicative of elevated homozygosity and inbreeding due to mating of closely related individuals. Self-fertilization can be a major source of inbreeding which elevates genomewide homozygosity and thus should also create long ROHs. While ROHs are frequently used to understand inbreeding in the context of conservation and selective breeding, as well as for consanguinity of populations and their demographic history, it remains unclear how ROH characteristics are altered by selfing and if this confounds expected signatures of inbreeding due to demographic change. Using simulations, we study the impact of the mode of reproduction and demographic history on ROHs. We apply random forests to identify unique characteristics of ROHs, indicative of different sources of inbreeding. We pinpoint distinct features of ROHs that can be used to better characterize the type of inbreeding the population was subjected to and to predict outcrossing rates and complex demographic histories. Using additional simulations and four empirical datasets, two from highly selfing species and two from mixed-maters, we predict the selfing rate and validate our estimations. We find that self-fertilization rates are successfully identified even with complex demography. Population genetic summary statistics improve algorithm accuracy particularly in the presence of additional inbreeding, e.g., from population bottlenecks. Our findings highlight the importance of ROHs in disentangling confounding factors related to various sources of inbreeding and demonstrate situations where such sources cannot be differentiated. Additionally, our random forest models provide a novel tool to the community for inferring selfing rates using genomic data.

## Introduction

Understanding the causes of inbreeding and its consequences on genetic diversity and fitness has been an ongoing discussion in the field of evolutionary genetics (Charlesworth and Charlesworth 1987; Keller and Waller 2002; Charlesworth and Willis 2009). Inbreeding can result from two disparate processes: self-fertilization (selfing), which is a biological property of an organism or its species (Lande and Schemske 1985), or demographic change reducing the population size, most often resulting from environmental or other external forces. Distinguishing the causes of inbreeding provides insight into two key aspects of individuals and populations – to what degree do individuals self-fertilize, and to what extent is the observed pattern of diversity a result of inbreeding due to past demography.

A common characteristic to assess inbreeding in populations is runs of homozygosity (ROHs). ROHs are continuous segments of increased homozygosity and can show where an individual has inherited the same haplotype from each parent, an effect known as autozygosity (Broman and Weber 1999). Comparing characteristics of ROHs among populations or individuals can reflect past demographic history (Ceballos *et al*. 2018). Even though ROHs have been commonly used to assess inbreeding in small populations or single individuals, the precise patterns of ROHs resulting from biparental inbreeding due to demography are not clear. How selfing changes ROH characteristics remains equally unexplored, apart from insights provided by Bennett (1953), who regarded selfing as a factor to increase homozygous tract lengths. Assessing the characteristics of ROHs resulting from interactions of demographic change and self-fertilization has not been studied to date, and may provide useful power to distinguish sources of inbreeding.

ROHs have a history in population genetics dating back to work by Fisher (1954), Bennett (1953) and Stam (1980). This research focused on the length of segments identical by descent and number of points delimiting such segments, called ‘junctions’. Broman and Weber (1999) also found long homozygous segments to be commonplace in human families, indicative of identity by descent. More recently, the advent of high-density genotype data has allowed for the association of long homozygous segments with consanguineous mating and human recessive disease (Woods *et al*. 2006; Gibson *et al*. 2006), at which point the term ‘ROH’ was coined (Lencz *et al*. 2007). ROHs have been analysed in the context livestock breeding (Peripolli *et al*. 2017; Doekes *et al*. 2019; Nosrati *et al*. 2021), demography of human populations (Pemberton *et al*. 2012; Gibson *et al*. 2006; Skourtanioti *et al*. 2023; Rivollat *et al*. 2022; Renaud *et al*. 2019; Lipson *et al*. 2022), recessive disease in humans (Woods *et al*. 2006; Lencz *et al*. 2007), inbreeding depression and genetic load (Szpiech *et al*. 2013; Bortoluzzi *et al*. 2020; Barragan *et al*. 2021), conservation genetics in small populations (Xue *et al*. 2015), and the recombination landscape (Bosse *et al*. 2012). Investigating ROHs in single individuals is particularly interesting in ancient DNA studies, where population-wide data is often not available (Skourtanioti *et al*. 2023; Rivollat *et al*. 2022; Renaud *et al*. 2019; Lipson *et al*. 2022). Though a large body of studies involving ROHs come from human and livestock genetics, the effect of selfing on ROHs has been neglected. Inbreeding, autozygosity and ROH statistics are undeniably related concepts, thus leveraging ROHs to distinguish sources of inbreeding has great potential.

Previous work has already shed some light on the characteristics of ROH segments expected under different demographic scenarios. Looking at counts and lengths of ROHs in human populations has shown different signatures due to recent consanguineous mating versus long-term shared ancestry (Kirin *et al*. 2010; Ceballos *et al*. 2018). Since recombination breaks up ROHs over generations, shorter segments reflect older relatedness. Given that length of ROHs is informative of past demography, analyses often categorize ROHs into bins to compare the frequency and distribution of different-sized ROHs (e.g., Pemberton *et al*. (2012); Skourtanioti *et al*. (2023)). Discretization simplifies the interpretation of continuous ROH data and allows one to capture non-linear relationships, but results in a loss of information and precision, and requires arbitrarily setting bin thresholds. One way to avoid these shortcomings is to compare the total proportion of the genome covered by ROHs (*F_ROH_*), which has also been found to be highly indicative of inbreeding (Keller *et al*. 2011). However, inbreeding may not be due only to demographic history, but also due to partial or full selfing (Lande and Schemske 1985).

Theory predicts that partial or full selfing increases the frequency of homozygous genotypes and decreases the effective recombination rate (Charlesworth and Wright 2001). These changes will likely increase the frequency and lengths of ROHs. Although Bennett (1953) addressed the effect of selfing on junction-formation, and Palamara *et al*. (2012) studied the relationship between identity by descent in a coalescent framework, the details of how patterns of ROHs are impacted by selfing, demography, or their interaction remain unknown.

The estimation of selfing rates is interesting for many biological questions, but historically required either extensive experimental setups, advanced knowledge the model system, or assumes that inbreeding is only due to selfing. Conventionally, the inference of selfing rates requires marker analysis of experimental progeny arrays to track the inheritance of a few independent loci (Ritland 2002; Koelling *et al*. 2012; Colicchio *et al*. 2020). The selfing rate (*σ*) can also be determined by using population genetic summary statistics. Assuming that inbreeding is only due to selfing, many studies use the *F*-statistics to estimate *σ* (*σ* = <u>^2*F*^</u>, Allard *et al*. (1968); Wright (1984)). Alternatively, *σ* can also be calculated based on linkage disequilibrium (LD) (Cutter 2006) or using an extension to a popular Bayesian Clustering approach (Gao *et al*. 2007). Another implementation to estimate the selfing rate uses identity disequilibria (ID) (David *et al*. 2007; Hardy 2016), a measure of association between homozygosity and/or heterozygosity across loci (Weir and Cockerham 1973). Tailored towards whole genome sequencing data, methods like eSMC and teSMC attempt to estimate selfing rate as well as the demographic history of the population based on the sequential Markovian coalescent (Sellinger *et al*. 2020; Strütt *et al*. 2023). While methods based on the *F*-static, LD or ID assume that the population is at equilibrium, which is often violated in natural populations (Brandvain and Wright 2016), other methods require knowledge of key parameters specific for the focal population or species. In this study we show that algorithmically capturing differences in ROH statistics by employing random forests provides a viable method to predict the selfing rate, even under complex demographic scenarios.

Random forests (Breiman 2001) are a machine learning tool for classification or regression and serve as a suitable approach to algorithmically model how highly dimensional predictor variables (’features’) are able to predict a target variable. This approach eliminates the need to bin ROHs and thus the subsequent loss of information from discretizing continuous data. Additionally, random forests can rank features according to their predictive power (’variable importance’). This makes random forests an ideal tool to explore how demography and selfing influence ROH characteristics.

In this study, we explore how demography and selfing shape patterns of ROHs. We use these ROH patterns and characteristics to distinguish sources of inbreeding to be able to infer the selfing rate in a population and a broad category of demographic history. By investigating the prevalence and variability of ROH characteristics in simulated populations, we identify ROH patterns specific to the type of inbreeding the population experienced. We find that mean lengths and counts of ROHs are indicative of past demographic change while the proportion of the genome covered by ROHs (*F_ROH_*) is influenced by the selfing rate. Using ROH features, we create random forest models to predict the selfing rate and demographic parameters. We then validate these predictions using additional simulations and test the algorithm in empirical data, confirming published differences in selfing rates among populations and taxa. We provide our pretrained, random forest machine learning models as a bioinformatic tool for predicting contemporary selfing rates from genomic data in natural populations. Our results furthermore give insight into how ROHs are shaped by different evolutionary forces, helping to disentangle inbreeding effects arising from selfing or demographic sources.

## Results

Using individual-based, forward-time simulations in SLiM v3.7.1 (Haller and Messer 2019), we modelled various demographic scenarios and selfing rates. We simulated scenarios of admixture, admixture followed by isolation, and several types of bottlenecks across a range of population sizes. As a null comparison we simulated cases of constant-size populations. Selfing rate was set within each simulation at a constant value, and we varied this rate from zero (obligate outcrossing) up to one (obligate selfing) across different replicates.

We initially focused on variation in counts of ROH and the proportion of the genome covered by ROHs (*F_ROH_*), since these have been previously described as indicative of differences in past and recent inbreeding (Kirin *et al*. 2010; Pemberton *et al*. 2012; Ceballos *et al*. 2018). In constant-size, outcrossing populations, i.e., populations without complex demographic or mating-type effects on inbreeding, we found that the *F_ROH_* is never greater than one half. *F_ROH_*decreased as *N* increased, except in the case of extremely small populations (*N* = 10) where *F_ROH_* ranged from 0 to 0.37 (maximum *F_ROH_* observed in any of our ideal, constant-size populations; Fig. 1, top-left panel). Mean counts of ROHs per individual consistently increased from smaller to larger *N*. Though it is not necessary in the simulations, in all cases counts were standardized by the chromosome length to be consistent with later analyses of empirical data across organisms of various chromosome numbers and sizes.

**Fig. 1:**
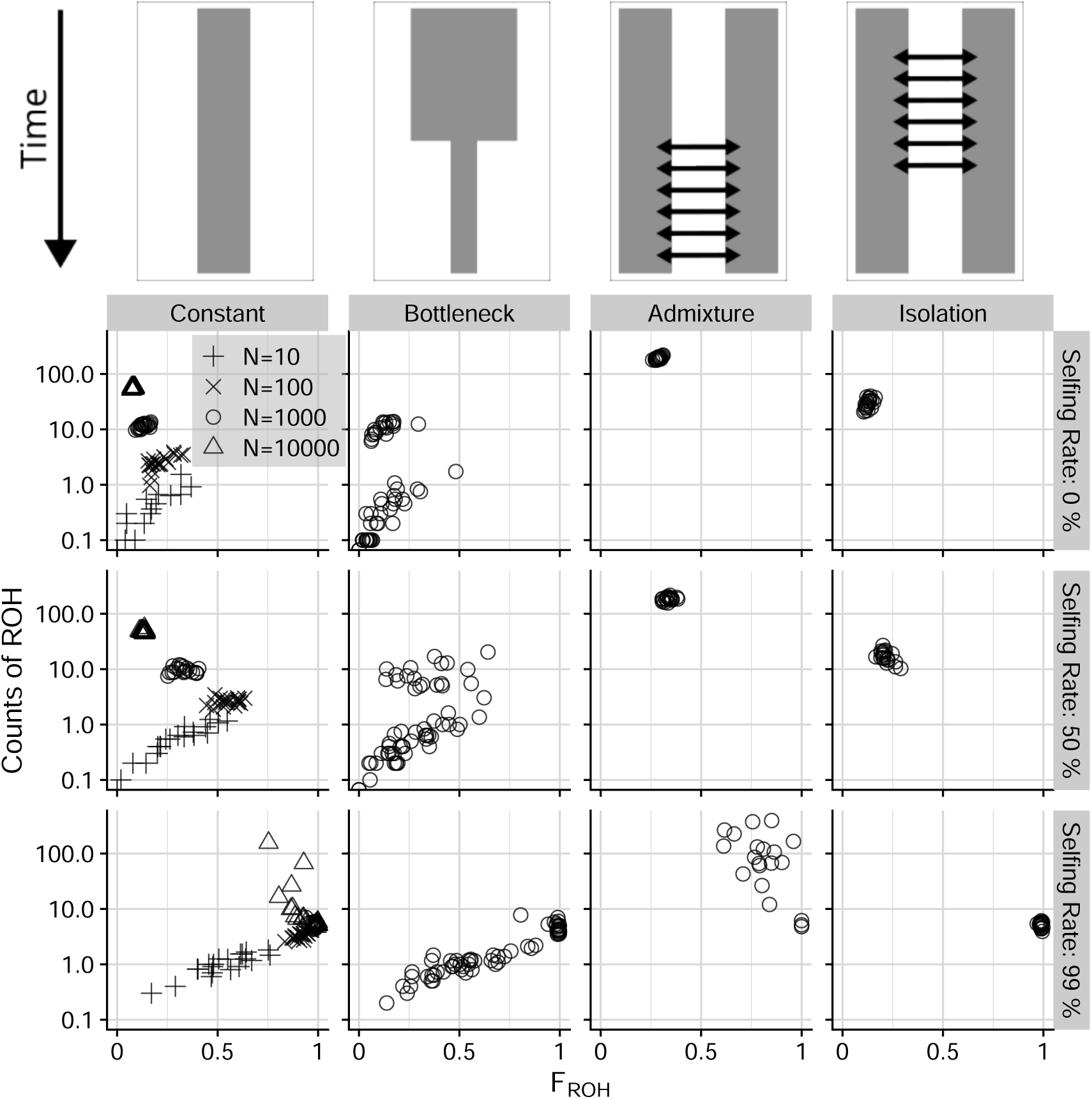
ROH patterns across various demographic histories and self-fertilization rates. Simulated populations either had a constant population size or underwent a demographic history of a bottleneck, admixture, or isolation after admixture. Each scenario was replicated at different selfing rates (obligate outcrossing, top row; 50% selfing, center row; 99% selfing, bottom row). Constant population size simulations show four values of *N*, while all other scenarios visualize only *N* = 1000 (open circles). Additional details distinguish the different types of bottlenecks are shown in Fig. S1 and Fig. S2.

Demography can drastically alter the patterns of ROHs. In comparison to constant-size, outcrossing populations, demography can have either no effect on ROH counts and *F_ROH_*, or conversely, result in substantially different ROH statistics (Fig. 1). For bottlenecked scenarios, ROH patterns differed depending on the type of bottleneck the population was exposed to, and on the initial population size (Fig. S1, S2). Some types of bottlenecks resulted in reduced *F_ROH_*, while others showed increased *F_ROH_*. Counts of ROHs were either unaffected or reduced. Our other two simulated demographic histories, those of admixture or admixture followed by a period of isolation showed increased ROH counts, and either increased *F_ROH_* (admixture) or unaltered *F_ROH_* (isolation).

We simulated three classes of bottlenecks, one with a gradual reduction and recovery in *N*, and two with instantaneous changes in *N* which was reduced by 1/100*N* for 5000 generations (long bottleneck) or 500 generations (short bottleneck). We set the initial population size for these three bottleneck types to *N* = 1000 or 10000, and documented ROH characteristics during the contracted period as well as after a period subsequent to the population size recovering. Bottlenecks resulting from a gradual reduction in population size did not greatly impact the distribution of *F_ROH_* and ROH counts. However, long and short bottlenecks with instantaneous size change had strong impacts on ROH characteristics. Long bottlenecks resulted in greatly reduced counts of ROH, decreased on average by 30-fold relative to constant size populations. Changes in *F_ROH_* depended on the initial population size. At *N* = 1000 *F_ROH_*changed only minimally on average, though had increased variance relative to constant size populations. For *N* = 10000 *F_ROH_* increased more than 2-fold, from 0.08 to 0.22. Even though this magnitude of change for *F_ROH_* was greater at a larger *N* and resulted in a higher *F_ROH_* for larger *N* bottlenecks, the range of values for *F_ROH_* was overlapping between both cases of *N* bottlenecks we simulated. Short bottlenecks resulted in substantially reduced counts in populations with small sizes (*N* = 1000, 40-fold decreased counts), as well as slightly decreased *F_ROH_*. The reduction in counts was less pronounced at bigger population sizes (*N* = 10000, 4-fold decrease), but increased *F_ROH_*. During recovery from a bottleneck, counts were still reduced compared to non-bottlenecked populations, but showed some recovery back towards higher counts for long and short bottlenecks only at *N* = 1000 (Fig. S1 left versus right column). *F_ROH_*reduced during bottleneck recovery, to a larger extent for bigger populations (*N* = 1000), bringing *F_ROH_* closer to levels observed in constant-size population cases.

Admixture created much higher counts of ROH per individual, increasing up to 17-fold. In *N* = 10 to 1000 scenarios, admixture also increased *F_ROH_* up to 2-fold. For *N* = 10000 admixture cases, *F_ROH_*was only slightly reduced. Historical admixture followed by a period of isolation led to reductions of counts in smaller populations (up to 2-fold, N=10 and 100), while bigger populations had increased counts (up to 2.5-fold, *N* = 1000 and 10000), therefore not as high as populations still experiencing admixture. *F_ROH_* in isolated populations was modestly decreased (N=10, 2-fold), without changes (*N* = 100 and 1000) or slightly increased (*N* = 10000).

In addition to demographic scenarios, we investigated the effect of self-fertilization on counts of ROH and *F_ROH_*. Increased selfing generally led to increased *F_ROH_* and reduced counts of ROH. Without complex demography, selfing increased *F_ROH_* by up to 12-fold, with 99% selfing causing *F_ROH_* values close to or equal to 1. Counts showed a less clear pattern, with overall more variance at each population size. Mean counts increased 3-fold for small populations (*N* = 10), while the pattern reversed in bigger populations, and resulted in counts reduced by 3-fold on average (*N* = 10000).

When selfing co-occurs with complex demographic history, changes to *F_ROH_* and ROH counts also interact, especially with more extreme selfing. *F_ROH_* still increased with higher selfing rates in all of our demographic scenarios (increase up to 12-fold for any bottleneck type, 11-fold for isolation and admixture). High selfing (*σ* = 0.99) in admixed populations resulted in both highly variable mean counts of ROH and *F_ROH_*. Counts ranged from values *>* 100 to *<* 10 and overlapped with all other demographic scenarios. Isolation with extreme selfing also overlapped with some cases in every other extreme selfing demographic scenario simulated.

Given the complex changes of ROHs to different sources of inbreeding, we compared statistics describing the distribution of ROHs in greater detail with other measures of inbreeding and selfing-rates. We calculated Pearson correlation coefficients between the classic *F*-statistic (*F*, inbreeding coefficient) and Tajima’s *D*, key indicators for inbreeding and demographic change, and ROH statistics calculated for every simulated population (median length, mean number per individual, mean *F_ROH_*, mean length of non-ROH segment, and variances of these). In all demographic scenarios, high selfing was associated with high *F_ROH_* (*R* = 0.79), but not counts (*R* = *−*0.13, Fig. S3). Median and variance of ROH lengths were also correlated with selfing (*R* = 0.57 and *R* = 0.58), and *F* showed the highest correlation (*R* = 0.93).

Median ROH length was correlated with *F_ROH_*, and to a lesser degree with its variance (*R* = 0.67 and 0.61, respectively). *F*, *F_ROH_*, length and length variance all showed positive correlation coefficients and clustered together according to the angular order of eigenvectors (Friendly 2002) (Fig. S3). Other variables were negatively correlated with inbreeding-related parameters, e.g., count of ROH, gap variance and Tajima’s *D*. Interestingly, Tajima’s *D* was correlated positively with count of ROH (*R* = 0.61).

The differences in associations of ROH characteristics with selfing and demography motivated us to explore if ROH statistics can help distinguish causes for inbreeding using machine learning. We used ROH variables as features to predict the selfing rate using random forests (Fig. 2A), a method that can capture complex, non-linear relationships in the data. From our simulations we used 80% of replicates for model training, and the remaining 20% as test data to assess predictive power of trained algorithms. We compared three different regression random forest models to predict the selfing rate using different variables as features: (1) a model using only ROH variables (we refer to this model as ‘RF-ROH’), (2) a model only using only *F* and Tajima’s *D* (’RF-stat’), and (3) a model using *F* and Tajima’s *D* in addition to ROH variables (’RF-full’). We found good performance and low error in the validation set for the RF-ROH model (*RMSE* = 0.114), but additional population genetic statistics improved the performance (RF-full, *RMSE* = 0.093, Fig. 2B, D). Only using population genetic summary statistics (RF-stat) also resulted in acceptable performance (*RMSE* = 0.111).

**Fig. 2:**
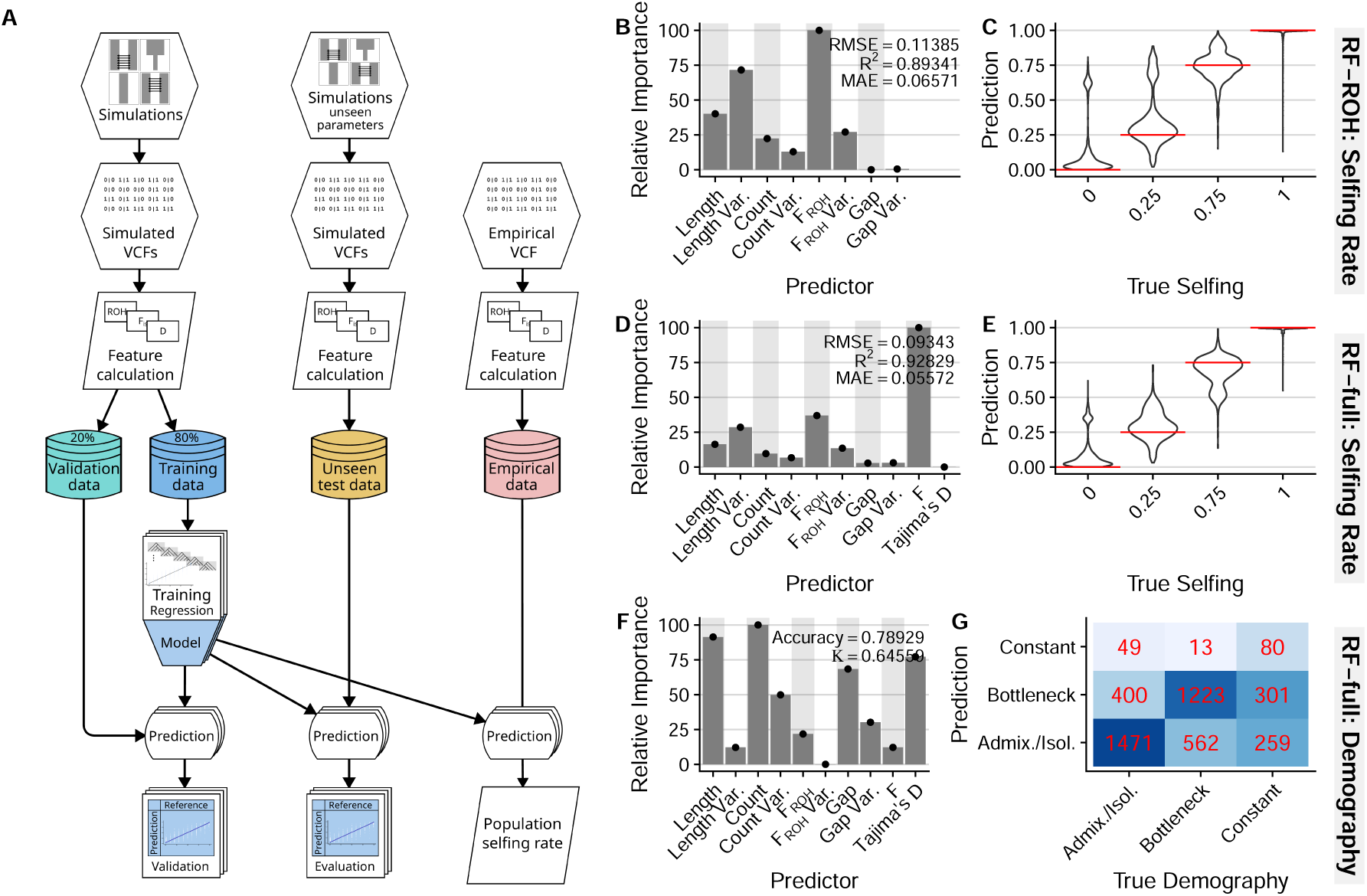
Outline of the random forest training and evaluation process (A). Relative feature importance and performance statistics for a model predicting selfing rates with features from ROH statistics only (validation data, B). We applied this model to data from simulations with novel, unseen input parameters (evaluation data) and predicted selfing rates (C). Violin plots show the distribution of predicted values for selfing rate, the red horizontal lines show true selfing rates. Relative feature importance and performance of a random forest model combining ROH and summary statistics (validation data, D) improved the predictions in the evaluation data (E). Relative feature importance (validation data, F) and confusion matrix evaluating classification prediction of demographic history from a random forest model using all features (evaluation data, G).

For more comprehensive testing of our previously trained models, we created an evaluation dataset with unseen simulation parameters, e.g., with mutations under selection as well as additional values of selfing rates (unseen test data in Fig. 2A). All models, RF-ROH, RF-full (Fig. 2C, E) and RF-stat were robust to unseen simulation parameters, albeit with some bias towards more intermediate selfing rates caused by selection (Fig. S4).

We compared our random forest models with a number of different models that predict selfing rate from *F* and ROHs to see if our approach provided any improved inference ability. A frequently used population genetic model predicts *σ* from 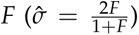 (Allard *et al*. 1968; Wright 1984). We also tested a linear regression model using *F* as a single predictor, and a LASSO regression with all predictors. The linear model has the advantage of being easily interpretable, while the LASSO regression allows for feature selection in models with high levels of multicollinearity. We the evaluated the performance of all models by comparing the RMSE (Fig. S5, S6). In both test and validation sets, random forest models using all predictors performed the best, with RF-full showing the best performance. In comparison to RF models, the conventionally used relationship between *F* and *σ* (Allard *et al*. 1968; Wright 1984) particularly underestimated in scenarios with lower selfing (*RMSE* = 0.181) and had the worst performance of all evaluated models. In the test set with unseen simulation parameters, despite elevated error, the RF-full model showed superior performance compared to other models (*RMSE* = 0.118, Fig. S6). Performance ranking based on RMSE did not change compared to validation set, except the RF-ROH model now performed worse compared to the simple linear regression with *F*. The LASSO model performed worse than the RF-stats model, while the population genetic model was biased toward low selfing rates. All non-RF models had the added disadvantage that they occasionally produced selfing rate estimates that fell outside of the biologically meaningful range (*<* 0 or *>* 1).

We also wanted to test if demographic scenarios can be predicted from ROH characteristics. To do this, we used random forest classification models and the same predictors as with selfing rate models. We trained on three broad demographic categories (constant, bottleneck, admixture), and evaluated the performance based on predictive accuracy. Accuracy was highest when the algorithm was trained on all predictors (*Accuracy* = 0.79 in validation; Fig. 2G, S7). We also tested classifying 28 specific demographic scenarios, described by a combination of the three broad demographic categories (constant, bottleneck, admixture), *N*, and specific characteristics within each demography (bottleneck types, contracted or recovered, admixed or isolated population, and all other combinations). These models performed worse compared to predicting broad categories (e.g., using all predictors, *Accuracy* = 0.64 in validation; Fig. S7). The model trained on all predictors again performed best. Predictions were less accurate for scenarios with smaller *N*, in particular for simulations with stable, constant size *N* (Fig. S8).

Random forests have the additional advantage to assess the relative importance of features, allowing us to better understand which ROH predictors are characteristic for the type of inbreeding the population was exposed to. We noticed the variable importance for ROH features to be context-dependent, where ROH count and length were relevant for demographic prediction, while for selfing rate *F_ROH_* and variance of length were highly meaningful (e.g., compare Fig. 2D with Fig. 2F, S7). Reassuringly, we recognized a similar heterogeneity when modeling only with *F* and Tajima’s *D*, whereby only *F* is important for selfing rate, and only *D* for demographic prediction.

To reinforce validity of the findings from variable importance in the random forest models, we conducted a principal component analysis (PCA) from the training set data and calculated loadings. The first two principal components both separated selfing rate and demography (Fig. 3A), and explained more than 50% of the variance (Fig. 3B). The first PC primarily explained variance in selfing rate, while the second PC explained more variance due to demographic differences. *F*, *F_ROH_*, ROH length and its variance all showed high loading on the first PC, and were roughly parallel to the selfing rate gradient, in concordance with the RF variable importance. We detected high collinearity for inbreeding coefficients with *F_ROH_*, ROH length and length variance, similar to findings from linear correlations (Fig. S3) Orthogonal to this was Tajima’s *D*, ROH count and its variance, gap variance, all loading onto the second PC. Although the first two axes helped to separate selfing rates and demographies, both effects still remained partially confounded.

**Fig. 3:**
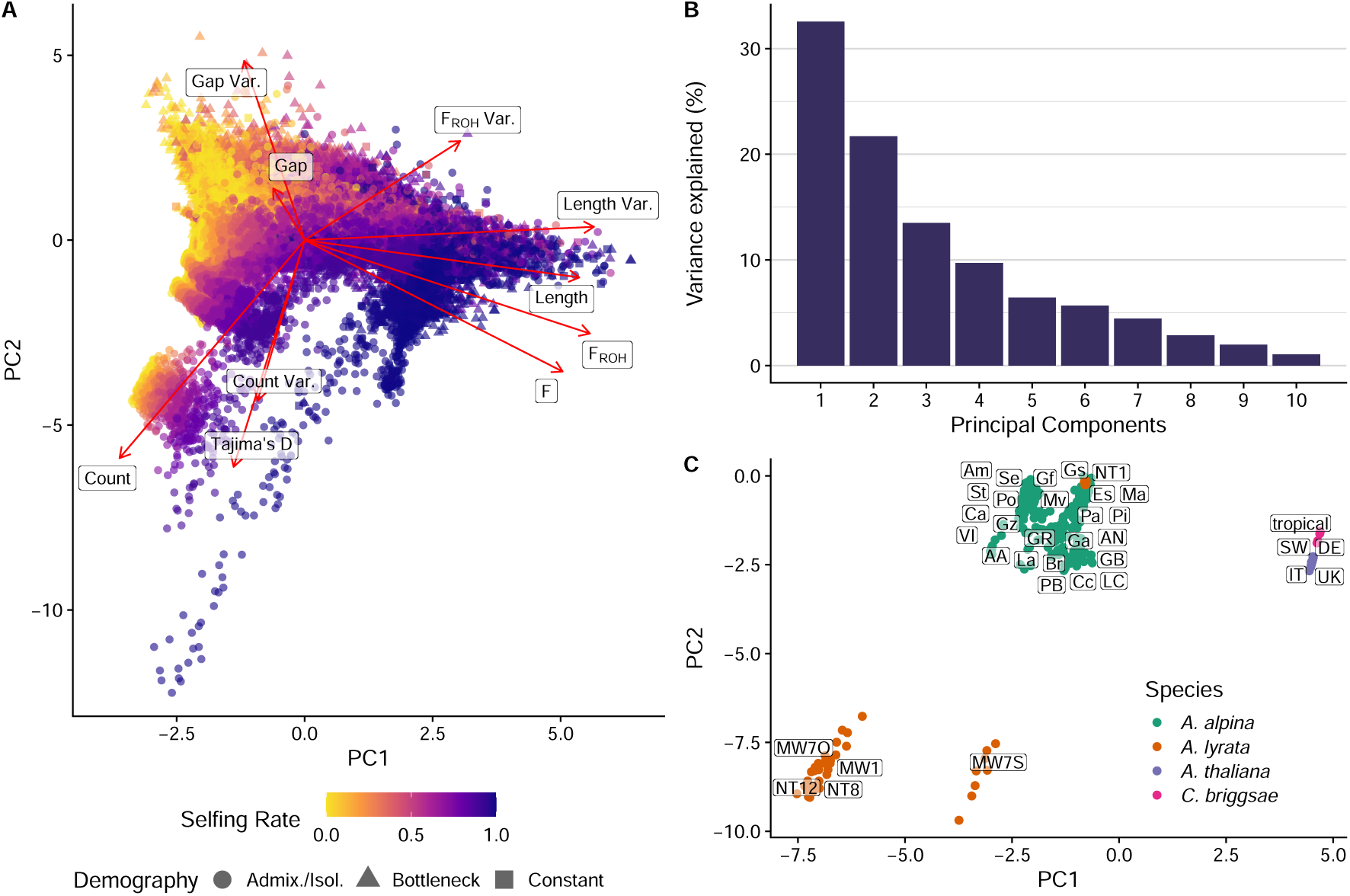
(A) A principal component analysis of all predictor variables and loadings using the simulation training set across all simulated selfing rates and demographic categories highlights the distinctions discernible based on different features. PC1 explained 32.6% and PC2 21.7% of the variance. (B) Scree plot for the PCA conducted in (A) highlights that the first two principal components capture *>* 50% of the variance explained. (C) PCA results for ROH features of chromosomes from empirical datasets of mixed mating *A. alpina* (green), *A. lyrata* (orange), and selfing *A. thaliana* (purple) and *C. briggsae* (pink), projected onto the same principal component space established in (A). Text labels indicate population names, for key see supplementary table S1.

We compared our findings to empirical data from four whole-genome sequencing datasets. We assessed mixed-mating species, with 24 populations of *Arabis alpina* (Laenen *et al*. 2018; Rogivue *et al*. 2019; Zeitler *et al*. 2023) and six populations of *Arabidopsis lyrata* (Kolesnikova *et al*. 2023), as well as two highly selfing species, with four populations from the model plant *Arabidopsis thaliana* (Alonso-Blanco *et al*. 2016) and one population of the nematode *Caenorhabditis briggsae* (Thomas *et al*. 2015). The *A. alpina* populations were originally sampled along a well documented expansion gradient, i.e., populations with fundamental demographic differences (Zeitler *et al*. 2023). We conducted a PCA on normalized ROH data, and mapped the empirical data into the same principal component space that was established for simulated data. Four outcrossing *A. lyrata* populations (NT12, NT8, MW1, MW7O) clustered together, while the two self-compatible populations (MW7S and NT1) did not (Fig. 3C). Self-compatible NT1 clustered closely together with self-compatible *A. alpina* populations from Switzerland (Ma, Es, Pi, Pa). Only focusing on PC1 and comparing the cluster of *A. alpina* populations, we found a gradient indicative of continuous variation in ROH data. This gradient shows a gradual shift from self-incompatible populations (Greek population VI and populations from the Italian peninsula, Am, Gs, Gz, Mv, AA, Ca, Gf, Po, Se, St) in positive direction on PC1 towards self-compatible Swiss, French, and Spanish populations (Pa, Pi, Es, Ma, GB, Ga, Cc, La, Br, PB, LC, AN). *A. thaliana* and *C. briggsae*, both highly selfing, clustered together and were separated from mixed mating *Arabis* and *Arabidopsis* populations. The second axis separated *A. lyrata* populations from the the other species and *A. lyrata* population NT1.

We analyzed ROH counts and *F_ROH_* in our empirical data and observed similar patterns of distinct clustering as in the PCA (Fig. 4A). NT1 and other selfing populations had higher *F_ROH_* compared to outcrossers, *A. lyrata* had higher counts than *A. alpina*. *F_ROH_* for *A. alpina* and *A. lyrata* fell within the same range, except NT1 which showed *F_ROH_* values similar to selfing *A. thaliana* and *C. briggsae*, near one. ROH counts of *A. thaliana* and *C. briggsae* were similar (mean ROH count per basepair and species 4 *×* 10*^−^*^8^ and 6 *×* 10*^−^*^8^, respectively), and much lower compared to mixed maters (1 *×* 10*^−^*^5^, 5 *×* 10*^−^*^5^ for *A. alpina* and *A. lyrata*). These observations broadly matched the patterns of simulated scenarios where high selfing shifts *F_ROH_* closer to one and counts to be relatively lower (Fig. 1).

**Fig. 4:**
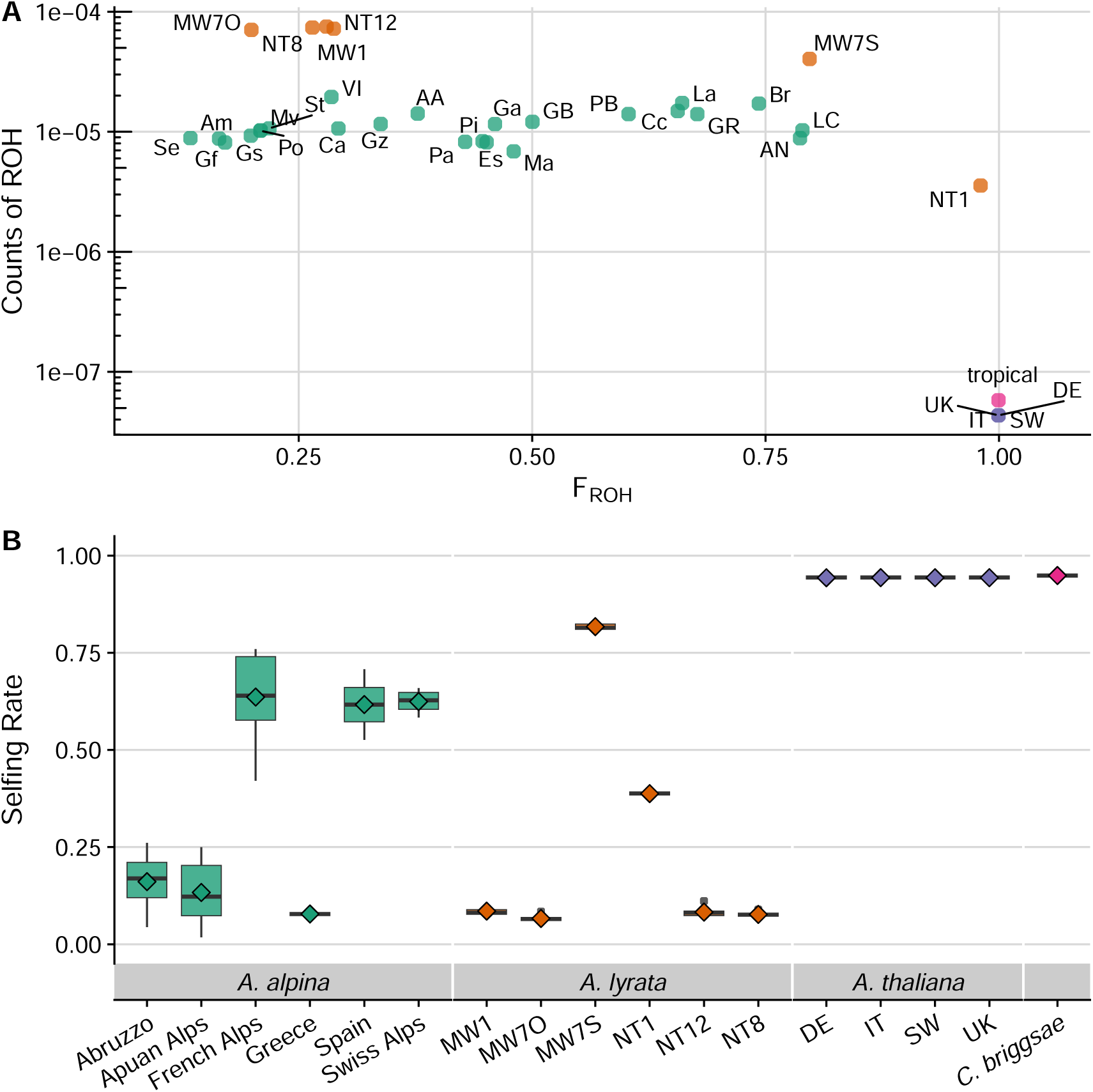
Average *F_ROH_* and average counts of ROHs averaged within empirical populations show distinct patterns of clustering (A). Using the sequential random forest model (see text), we predicted selfing rates for these populations. These species span mixed mating systems, *A. alpina* and *A. lyrata*, as well as known highly selfing model organisms, *A. thaliana* and *C. briggsae* (B). Each box-and-whisker plot represents the distribution of estimates of selfing across chromosomes either within regions (of several subpopulations, *A. alpina*) or within populations (*A. lyrata, A. thaliana, C. briggsae*). Diamonds indicate group means.

Most relevant for future applications of our RF algorithm, we inferred selfing rates in these empirical populations. We first used the RF-full model, which resulted in estimates of selfing rate ranging from 0.07 to 0.85 (populations from *A. alpina* and *C. briggsae*, respectively; Fig. S9). Given the surprisingly low estimates for highly selfing species, we attempted another set of inferences by first estimating *N* in a RF classification model and then incorporated this estimate as a feature in a second RF regression model to predict selfing. This sequential modeling approach (elsewhere in this manuscript referred to as the ‘sequential model’) performed well (Fig. S10, S11), similar to the RF-full model.

This prediction increased the mean estimates selfing from 0.83 to 0.94 for *A. thaliana* and from 0.86 to 0.95 for *C. briggsae*. We therefore proceeded with the sequential model for the prediction of selfing rates in the remaining mixed-mating species. We estimated selfing rates less than 0.16 for populations thought to be self-incompatible and between 0.39 to 0.82 for populations described as self-compatible (Fig. 3B). The self-incompatible population from Greece (VI) had the lowest selfing rate (0.08) and putatively outcrossing Italian populations had moderately low selfing rates (0.16, 0.13 for Abruzzo and Apuan Alps).

## Discussion

We have shown that different sources of inbreeding shape patterns of runs of homozygosity in the genome in different manners. These patterns thus allow us to identify contemporary selfing rates and demographic histories even when they simultaneously cause (or prevent) inbreeding. We present a machine learning algorithm that can serve as a method to infer selfing rates in natural populations, and compare it to several others models for predicting the selfing rate from ROH data.

ROHs provide unique insights into mating system and demography, and have been frequently used to quantify and compare the levels of inbreeding within and among populations. Several statistics can summarize ROHs and allow for the comparison of patterns to infer the respective evolutionary processes generating them. Ceballos *et al*. (2018) commented that the number of ROHs within an individual’s genome is mostly influenced by demographic change the population is experiencing. Given these predictions, we expected fewer ROHs in populations with larger ancestral effective population sizes, as well as more and longer ROHs in smaller populations. Our results confirm this and show that longer ROHs are a signal of more recently shared common ancestry and inbreeding, e.g., in small populations. Inbreeding due to demography was also identifiable by differences in counts, which served as a useful predictor, being more prevalent in larger populations.

Bottlenecks had ROH patterns similar to smaller populations, reflecting similar effective population sizes. ROH patterns of bottlenecks were also dependent on differences in timing during or after bottleneck exposure. We found that counts of ROHs and *F_ROH_* changed depending on the type of bottleneck, indicative of differences in time passed since reduction in population size. Recovery from bottlenecks resulted in increased but not re-equilibrated ROH statistics. These findings are consistent with theory, predicting that the recovery of heterozygosity after a bottleneck is dependent on the number of generations (Nei *et al*. 1975; Allendorf 1986). While populations recovering from inbreeding events showed increased heterozygosity and had ROHs subdivided into smaller segments, smaller populations did not show overall reductions in homozygosity. Overall, demographic history that impacts population size over different time spans clearly led to different ROH patterns.

Admixture created one of the most distinguishable patterns of ROHs in our simulations. Contrary to the prediction by (Ceballos *et al*. 2018), we found that admixture results in increased numbers of ROHs. This is because recombination effectively partitions ROHs into more but smaller segments. *F_ROH_* can still remain similar or moderately higher than constant-size populations. Interestingly, the moderate increase in *F_ROH_*may also occur with a Wahlund effect, where a departure from Hardy-Weinberg equilibrium due to population subdivision is observed and results in a higher *F*-statistic. The rate of migration between two admixing populations is also likely to impact the patterns of ROHs observed.

Selfing is known to increase homozygosity and reduce recombination at the population level, and therefore resulted, as expected, in longer ROHs and increased *F_ROH_* relative to outcrossers. *F_ROH_* close to one indicated high self-fertilization, in line with expectations of increased *F*-statistics due to selfing. These patterns hold regardless of the underlying demography, whether constant size, bottlenecks, or isolation. However, ROH statistics from admixed populations are outliers in terms of their interactions with selfing. Under high selfing rates, both means and variances of *F_ROH_* and counts of ROH increased. We hypothesize that these patterns are created by rare outcrossing events in otherwise selfing individuals, resulting in gene flow and sporadic introduction of new haplotypes. The high variance in ROH statistics thus represents differences in age and frequency of such gene flow events. Furthermore, differences in selfing rate among admixing populations may also create new patterns we have not yet explored, particularly if highly selfing populations mix with largely outcrossing populations.

The unique signatures of ROHs associated with different demographies or selfing rates provided useful power for our machine learning approach to distinguish causes of inbreeding. Ranking the importance of ROH parameters helped us to better understand these distinguishing characteristics. We find that ROH counts predict type of demography the population experienced best, and *F_ROH_* helps distinguish outcrossing from selfing. Results from PCA loadings were in agreement with these findings and highlighted the importance of ROH lengths for demographic inference. Tajima’s *D* reflected changes due to demographic history, as expected from theory (Tajima 1989). However, when predicting demographic scenarios using ROHs, we find *F_ROH_* to be less helpful, contrary to what is suggested in existing literature (Keller *et al*. 2011; Peripolli *et al*. 2017). *F_ROH_* is instead more informative to quantify the degree of self-fertilization. When analyzing ROHs in a system that does not self, it may thus be more helpful to compare mean counts and lengths of ROHs among populations reflecting bi-parental inbreeding. The other conventional population genetic summary statistic we used, the inbreeding coefficient *F*, is also relevant for inferring the selfing rate. The high importance of *F* and *F_ROH_*for estimating selfing rate is reassuring, considering the conventional approach to estimate selfing rates uses *F*-statistics.

### Empirical applications of random forest inference

Given that random forest models were robust to unseen population genetic parameters, we proceeded with predictions from empirical genomic data. For the highly selfing species *A. thaliana* and *C. briggsae* we expected rates of approximately *>* 0.98 (experimentally estimated in outcrossing rate, Snape and Lawrence (1971); Redei (1975)) and *>* 0.9989 (genetically effective outcrossing rate, Thomas *et al*. (2015)), respectively. The model predicting selfing rate from all features without taking into account the population size (RF-full) produced lower-than-expected selfing rates for species known to be highly selfing. We thus created a second model, predicting selfing in bins of the predicted census size. Even though model performance was not noticeably higher in tests with unseen simulated data, we found that the sequential model estimates higher, more realistic selfing rates for *C. briggsae* and *A. thaliana*. Although no data is available for experimentally estimated outcrossing rates our focal populations, *A. thaliana* is known to be highly selfing (Redei 1975), However, in the wild, outcrossing rates may be slightly higher (up to 8 %) compared to greenhouse conditions (Snape and Lawrence 1971). Expanding our tests beyond plants, our model predicted that *C. briggsae* may occasionally outcross, even though the entire sequenced genome is covered by ROHs (*F_ROH_* = 1). Whether this is an issue in our model or a realistic feature of the species is yet to be determined. *C. briggsae*, similar to *C. elegans* is described as highly selfing, but analyses based on LD, the effective recombination rate, and the effective population size (*N_e_*) suggest that occasional outcrossing is possible (Cutter 2006; Thomas *et al*. 2015). The literature describes outcrossing rates as very low (*∼* 10*^−^*^3^), however, these rates are effective outcrossing rates based on the *F*-statistics and are thus also influenced by joint effects of non-random mating and outcrossing, while our rates should correspond closer to *in situ* outcrossing rates. Differences between the RF-full and the sequential model were only realized in these highly selfing species and are potentially indicative of other unknown biological factors that our simulations did not capture.

Predictions of selfing rates in populations with documented interactions of mixed mating and complex demographic confirmed a wide range of selfing estimates for different populations. Particularly in *A. alpina*, ROH characteristics predicting selfing rate showed considerable variation, potentially stemming from a higher variance in the true selfing rate or potential gene flow from adjacent populations. Complex history of a range expansion in these populations may also contribute to variability in ROH patterns (Laenen *et al*. 2018; Zeitler *et al*. 2023). Overall, selfing rates of *A. alpina* corresponded with presumed variation of self-compatibility described in the literature (Ansell *et al*. 2008; Tedder *et al*. 2011). Even though we only inferred selfing rates at a population level, there may also be variation within populations, as exemplified by estimates of inbreeding coefficients in *A. alpina* individuals (Zeitler *et al*. 2023). Studies comparing selfing rates with pollinator essays or progeny arrays would be useful to confirm the accuracy of our predictions and which level of population structure is most relevant to infer the selfing rate.

In *A. lyrata*, we found two mixed mating populations (NT1, MW7S) matching published populations that gained self-compatibility (Kolesnikova *et al*. 2023). Our results show that self-compatible NT1 is a mixed mater with outcrossing still occurring, while MW7S showed higher degrees of selfing. These selfcompatible lineages with intermediate selfing rates would allow for frequent admixture events, consistent with the analyses of Kolesnikova *et al*. (2023), which detected a considerable amount of contemporary gene flow between selfing and outcrossing populations.

Compared to other methods available for inferring the selfing rate from whole genome data, our method does not require previous knowledge of recombination or mutation rates, or a dedicated sampling strategy such as progeny-arrays. Our method is applicable to a wide array of natural populations because we trained on many different potential parameters, learning from variables influencing homozygosity and inbreeding. While we cover a broad range of common demographic scenarios, some non-random mating patterns, e.g., in experimental or breeding populations, or asexual reproduction will be unaccounted for. Some ROH features caused by selfing or demography may remain indistinguishable, e.g., small, highly selfing populations could be confused with bottlenecked ones, making demographic predictions of small selfing populations challenging (Fig. S8). Furthermore, some naturally occurring mutational and chromosomal properties were not factored into our simulations. Large scale chromosomal rearrangements and differences in interactions of these with demographic changes may lead to erroneous estimates. Variation of the recombination rate along chromosomes, such as hotspots or regions of reduced recombination will likely change how quickly unique ROH patterns vanish over time. This will in turn elevate the variance of ROH variables and could make predictions less precise. Another factor altering the prediction of selfing rates is selection. In our training set, all mutations were simulated as neutral, which resulted in biased predictions towards intermediate selfing rates. For high selfers, this bias could be caused by residual heterozygosity at sites with lethal mutations, which would split ROHs into more and smaller runs. The length of ROHs or *F_ROH_*may also be altered by mutations and subsequent selection. Selective sweeps will increase homozygosity and perhaps bias prediction, causing an outcrossing population with longer ROHs than expected under neutrality. Such artifacts could, and should, be circumvented by using only putative neutral loci, as is often done for inference methods biased by selection (Gutenkunst *et al*. 2009).

Our models estimate rates for single chromosomes and per population, even though in a biological sense, the outcrossing rate is a property of an individual. The rationale for estimating per-chromosome and population values is primarily a choice of computational efficiency but the approach also strengthens the algorithm against individual variation in sequencing quality resulting in heterozygosity outliers. To better understand the level of uncertainty when applying our models to estimate the selfing rate, it may be helpful to examine the range of values across various chromosomes. Some degree of uncertainty may still remain with the estimates. Future research analysing ROHs using our models could be able to distinguish between types of inbreeding and characterize *in-situ* outcrossing rates to make comparisons with model predictions. Such analyses would also allow for comparisons with environmental, population genetic and ecological variables influencing the selfing rate.

Predictive uncertainty may also arise from empirical sequencing data, as the number of accessible bases is often not same as the maximum length of a ROH, as some parts of the genome will not be covered by reads. Such areas may include centromeric and telomeric regions, because of read mapping difficulties in repetitive sequences. Technical difficulties may additionally cause biases in some sequencing datasets, as calling heterozygous genotypes is generally associated with higher error, but crucial for correct ROH inferences. Especially in highly selfing organisms, false heterozygous calls may cause incorrect predictions because long ROHs are interrupted, but the extent of this effect remains to be tested. Issues regarding heterozygosity and recombination rate can be mitigated by the user fine-tuning bcftools ROH to account for specifics in the focal data, e.g., by supplying a genetic map to the algorithm, running the program in genotype-likelihood mode, and fine-tuning transition proabilities. Another integral step before analysing selfing rates using our models should also be quality control of raw ROH data, and, if applicable, masking genomic regions where high error is suspected, in both ROH data and the reference genome. Accounting for erroneous heterozygosity in repetitive regions and selfing organisms, as well as extending the algorithm to single individuals to identify intra-population variation of ROHs, is a potential avenue for future improvement.

## Conclusions

ROHs provide unique insights into mating system and demographic history of populations. While the association between ROH length and bi-parental inbreeding is documented for empirical systems, our analysis connects specific ROH features with inbreeding-related properties of populations. We find that the proportions of ROHs, *F_ROH_*, are indicative of the degree of inbreeding by self-fertilization, while count and mean length of ROHs are associated with demographic parameters. Training machine learning models on such features allowed us to build algorithms able to predict the selfing rate and demographic history. This shows that ROHs capture useful genetic signatures in populations that distinguish inbreeding sources. Distinguishing sources of inbreeding contributes to a better understanding of the evolution of mating systems and how this may relate to demographic changes in populations. Our insights open avenues for future work to investigate the interaction between selection and ROHs, as such regions of low diversity harbour genetic load or may influence the potential to adapt to changing future environments.

## Material and Methods

### Simulations

Using SLiM 3.7.1 (Haller and Messer 2019), we simulated a range of demographic scenarios and selfing rates to understand how these parameters influence patterns of ROH statistics. These scenarios modeled populations of constant size, bottlenecks, or a history of admixture with or without subsequent isolation (Fig. 1). For constant size and admixture cases, population size was set to *N* = 10, *N* = 100, *N* = 1000 or *N* = 10000. Bottleneck scenarios were set to *N* = 1000 or *N* = 10000 and experienced a 100-fold instantaneous reduction in population size for 500 (“short bottleneck”) or 5000 (“long bottleneck”) generations, or a gradually decreasing population size for 2500 generations (“gradual bottleneck”). These demographic changes were initiated after a burn-in phase of 50000 generations. Admixture scenarios consisted of two subpopulations of equal size *N* and equal selfing rate. Migration between subpopulations began at 50000 generations with migration rate *m* = 0.1, and samples were either drawn 500 generations later (“early admixture”) or 5000 generations later (“admixture”). In the admixture scenario with subsequent isolation, we simulated 500 generations of admixture, followed by 4500 generations without migration (“isolation”). We set a constant selfing rate in any given simulated scenario, and varied this rate across different simulations from *σ* = 0 (obligate outcrossing, no incidental selfing), 0.01, 0.1, 0.2, 0.3, 0.4, 0.5, 0.6, 0.7, 0.8, 0.9, 0.99 or 1 (obligate selfing). Each individual had a diploid genome with a genome size of 1 *×* 10^7^ base pairs (bp) and a constant recombination rate of 1 *×* 10*^−^*^8^ per bp per generation. We simulated neutral mutations at a per base pair mutation rate of 7 *×* 10*^−^*^8^, and further simulations described below included selected variants. We output of a maximum of 100 randomly sampled individuals from each simulation, recorded at the end of each simulated scenario. For bottleneck simulations we additionally sample during the bottleneck contraction phase, 500 generations (’short’, ‘gradual’ types) or 2500 generations after the start of the contraction. In some bottleneck cases, due to the small population size, the number of individuals for contracted populations was less than 100. Every scenario was replicated 100 times.

For a second set of simulations to later assess “unseen” patterns from novel situations, we repeated the simulations described above but changed the initial parameter space to contain intermediate population size parameters, added a modifier that allowed for non-neutral mutations, and simulated two additional selfing rates. We simulated *N* = 5000 and *N* = 1000 for any demographic scenario, and additionally *N* = 500 and *N* = 50 for admixture and constant *N* settings. Selfing rate *σ* we set to 0, 0.25, 0.75 or 1. In addition to neutral simulations, these scenarios allowed for natural selection. Selected mutants occurred at the same rate of mutation per base pair; we simulated neutral, beneficial, deleterious, and lethal mutations, which occurred in proportions of 0.25, 0.001, 0.649, and 0.1, respectively. Lethal alleles had a selection coefficient of 1, and selection coefficients for deleterious and beneficial mutations were derived using an exponential function with a mean of -0.001 and 0.01, respectively. For beneficial and deleterious alleles, the dominance coefficients were set to h = 0.3, and for lethal mutations, to h = 0.02. We repeated this second set of simulations over 20 replicates.

### ROH statistics

We used bcftools roh 1.18 (Danecek *et al*. 2021; Narasimhan *et al*. 2016) to identify ROHs from our simulated results, which allows calculation of ROH either with genotype calls or genotype likelihoods and uses a hidden Markov model (HMM) to identify ROHs. We calculated ROH over the entire genome of every saved individual and then summarized the raw output over entire populations by calculating several summary statistics describing characteristics of ROH. For each population we counted number and median length of ROHs, the total proportion of the genome covered by ROHs (*F_ROH_*, Keller *et al*. (2011)), and the median length of non-ROH segments (“gaps”). In addition, we calculated the variances of these statistics, enabling us to describe the distribution of ROHs in each population with eight variables (supplementary table S2). To accompany these variables with more classical population genetic summary statistics to detect demographic change and inbreeding, we calculated inbreeding coefficient (*F*) estimates as a indication for non-random mating, and Tajima’s *D* (Tajima 1989), which can be used to detect demographic expansion or contraction. We calculated *F* using plink 1.90 (Chang *et al*. 2015) (–het) per individual, and Tajima’s *D* using vcftools 0.1.15 (Danecek *et al*. 2011) and summarized both for entire populations.

### Statistical Modeling using ROH Features

To test the suitability of ROH statistics for predicting selfing rates of populations with different demographic histories, we used random forest (RF) models. We used R 4.2.2 (R Core Team 2022), caret (Kuhn 2008), and the ranger implementation of random forests (Wright and Ziegler 2017). We divided the whole simulation dataset into three subsets. We split the primary dataset into training and validation sets, by assigning the first 80 replicates of every parameter combination to the first, and the remaining 20 to the latter. The supplementary simulation set with partially modified parameters was used as a test set, to verify model predictions in scenarios with unseen simulation parameters. We used a stepwise approach to allow the model training to use only ROH-statistics (“RF-ROH”), only population genetic summary statistics (“RF-stats”, *F* and Tajima’s *D*), or all population statistics (“RF-full”) as predictors. We additionally trained a model that first classified *N*, the census population size of at the sampling time point in classes of 10, 100, 1000 or 10000. Knowing *N*, we subsequently trained a regression RF with *N* as additional feature (“sequential RF”). For training *N* was known from the data, and during evaluation, *N* was inferred using the classifier model. We used all features for this sequential RF. First, we preprocessed features by standardization (centering, scaling). We additionally partitioned the training data into subsets using a grouped k-fold cross-validation strategy. During the training process, the number of folds matched the number of grouped k-folds and a grid search was employed to optimize hyperparameters. We estimated variable importance using a permutation approach and the root mean square error (RMSE) as primary performance evaluation metric.

Using the same data, we also trained classification random forest models to distinguish between 28 demographic categories. These target categories were combinations of identifiers for specific demography type (bottleneck types, admixture, isolation, constant) and *N*. Alternatively, we trained classifier models to predict more broadly between bottleneck, admixture/isolation, or constant size demographic categories. For these classifier models, we used the classification accuracy to evaluate performance. Classification accuracy is the proportion of correctly classified predictions in the dataset divided by the total number of instances.

To compare the selfing rate predictions from RF models with alternative approaches, we also used a regularized generalized linear model (Lasso regression, ‘LASSO-GLMNET’, Friedman *et al*. (2010)), two linear regression models and the frequently used formula 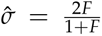 Allard *et al*. (1968); Wright (1984).

For the Lasso regression we used stratified bootstrap resampling with hyperparameter tuning for the regularization parameter to control model complexity and mitigate overfitting risks. We constructed a final Lasso regression model by selecting the penalty value that leads to the best performance based on RMSE from the hyperparameter tuning results. We also used a simple linear model to test an easily interpretable prediction of selfing rate *σ_F_* = *aF* + *b* with *F* as mean inbreeding coefficient. We applied the models in the test and validation sets and compared the results based on RMSE, mean absolute error (MAE) and *R*^2^ values. We also conducted a principal component analysis (PCA) and calculated loadings to further investigate the relationship of ROH, *F* and Tajima’s *D*. Regression and PCA modelling was conducted using the package tidymodels (Kuhn and Wickham 2020) and the comparison of *R* for features plotted using corrplot (Wei and Simko 2021).

### Empirical Datasets

To compare the results from simulations with empricial data from natural populations, we used published short read sequenced genomes of the mixed mating plants *Arabidopsis lyrata* and *Arabis alpina*, and data from the selfing model species *Arabidopsis thaliana* and *Caenorhabditis briggsae*. For *A. alpina*, we aquired data from published resources (Laenen *et al*. 2018; Rogivue *et al*. 2019; Zeitler *et al*. 2023), and followed previously published procedures to obtain a variant call set (Zeitler *et al*. 2023). For *A. lyrata* we used the same biallelic SNPs as described in the demographic modeling section of Kolesnikova *et al*. (2023) kindly provided by the authors. For *A. thaliana* we downloaded data from the 1001 genomes project website (https://1001genomes.org/data/GMIMPI/releases/v3.1/1001genomes_snpeff_v3.1/, (Alonso-Blanco *et al*. 2016)) and subsetted individuals sampled from four sampling locations, representing a wide geographical range (Germany: ‘DE’, United Kingdom: ‘UK’, Italy: ‘IT’, Sweden: ‘SW’). A VCF file containing variants for tropical nematode samples of *C. briggsae* was kindly provided by the authors; information on these individuals is available in the original publication (Thomas *et al*. 2015). As per the methods described in Thomas *et al*. 2015, genotypes in this last dataset were originally called as haploid, we converted these calls into diploids using a custom script. Using vcftools (Danecek *et al*. 2011) and bcftools (Danecek *et al*. 2021) we filtered all datasets to only retain sites with a maximum missingness of 20% and only biallelic sites. As with the simulations, we calculated ROH using bcftools roh, however, with empirical data we used the program’s ability to process allele counts from ‘AC’ and ‘AN’ annotations in the file, previously processed using freebayes 1.3.2 (Garrison and Marth 2012) and GATK SelectVariants 4.2.0.0 (Van der Auwera *et al*. 2013). Again, we calculated *F* and Tajima’s *D* using plink and vcftools (Chang *et al*. 2015; Danecek *et al*. 2011).

We then wrote a Rscript wrapper to apply *F*, Tajima’s *D* and ROH output from any population to estimate selfing rates (https://github.com/LZeitler/roh-selfing). We supply pre-trained models for RF-stats, RF-ROH, RF-full and sequential-RF to infer the selfing rate of any population with at least two individuals. The program calculates ROH statistics per basepair by dividing length, count, proportion and gap by the length of the chromosome, to account for differences in chromosome lengths between training and empirical sets. Accordingly, the program outputs selfing rate estimates for each chromosome. We applied the pre-trained models to ROH and summary statistics from our four sequencing datasets to obtain selfing-rates of populations with either variable selfing-rates documented in the literature (supplementary table S1, see also (Ansell *et al*. 2008; Tedder *et al*. 2011; Buehler *et al*. 2012) for *A. alpina* and (Kolesnikova *et al*. 2023) for *A. lyrata*) or as a empirical control of highly selfing organisms *A. thaliana* and *C. briggsae*.

## Data and code availability

Code, simulation output and pretrained models are available on GitHub (https://github.com/LZeitler/ roh-selfing).

## Acknowledgments

We thank Stephan Peischl for statistical advice, Polina Novikova and Alison Scott for providing access to the *A. lyrata* dataset, and Tyler Kent and Marinela Dukić for critical feedback on the manuscript.

Computation was performed in part on UBELIX (http://www.id.unibe.ch/hpc), the HPC cluster at the University of Bern. This research was funded by Swiss National Science Foundation Ambizione grant #PZ00P3_185952 to KJG.

## Author Contributions

## Supplementary Material

**Table S1:** Empirical populations with abbreviations, assumed mating and reference for sequences.

| Species | Population | Original name | Assumed mating | Reference |
| --- | --- | --- | --- | --- |
| *Arabidopsis lyrata* | MW7O | MW0079579,<br>MW0079581 | SI | Kolesnikova *et al.* (2023) |
|  | MW7S | MW0079456,<br>MW0079570 | SC | Kolesnikova *et al.* (2023) |
|  | MW1 | MW0158706,<br>MW0158707 | SI | Kolesnikova *et al.* (2023) |
|  | NT1 | NT1_1_1, NT1_2_1,<br>NT1_3_1 | SC | Kolesnikova *et al.* (2023) |
|  | NT12 | NT12_1_1, NT12_2_1 | SI | Kolesnikova *et al.* (2023) |
|  | NT8 | NT8_1_1, NT8_2_1,<br>NT8_3_1, NT8_4_1 | SI | Kolesnikova *et al.* (2023) |
| *Arabis alpina* | AA | no change | outcrossing | Laenen *et al.* (2018) |
|  | Am | no change | outcrossing | Zeitler *et al.* (2023) |
|  | AN | no change | selfing | Laenen *et al.* (2018) |
|  | Br | no change | selfing | Zeitler *et al.* (2023) |
|  | Ca | no change | outcrossing | Zeitler *et al.* (2023) |
|  | Cc | no change | selfing | Zeitler *et al.* (2023) |
|  | Es | no change | selfing | Rogivue *et al.* (2019) |
|  | Ga | no change | selfing | Zeitler *et al.* (2023) |
|  | GB | no change | selfing | Laenen *et al.* (2018) |
|  | Gf | no change | outcrossing | Zeitler *et al.* (2023) |
|  | GR | no change | selfing | Laenen *et al.* (2018) |
|  | Gs | no change | outcrossing | Zeitler *et al.* (2023) |
|  | Gz | no change | outcrossing | Zeitler *et al.* (2023) |
|  | La | no change | selfing | Zeitler *et al.* (2023) |
|  | LC | no change | selfing | Laenen *et al.* (2018) |
|  | Ma | no change | selfing | Rogivue *et al.* (2019) |
|  | Mv | no change | outcrossing | Zeitler *et al.* (2023) |
|  | Pa | no change | selfing | Rogivue *et al.* (2019) |
|  | PB | no change | selfing | Laenen *et al.* (2018) |
|  | Pi | no change | selfing | Rogivue *et al.* (2019) |
|  | Po | no change | outcrossing | Zeitler *et al.* (2023) |
|  | Se | no change | outcrossing | Zeitler *et al.* (2023) |
|  | St | no change | outcrossing | Zeitler *et al.* (2023) |
|  | VI | no change | selfing | Laenen *et al.* (2018) |
| *Arabidopsis thaliana* | DE | 9785, 9786, 9815,<br>9797, 9798, 9800, 9801 | selfing | Alonso-Blanco *et al.* (2016) |
|  | IT | 9673, 9672, 9671,<br>9670, 9669, 9668,<br>9973, 9667, 9666,<br>9665, 9664 | selfing | Alonso-Blanco *et al.* (2016) |
|  | SW | 9801, 9800, 9798,<br>9797, 9815, 9786, 9785 | selfing | Alonso-Blanco *et al.* (2016) |
|  | UK | 5276, 5279, 5726, 7064 | selfing | Alonso-Blanco *et al.* (2016) |
| *Caenorhabditis briggsae* | tropical | no change | selfing | Thomas *et al.* (2015) |

**Table S2:** Description of predictor variables derived from ROH statistics, and output variables.

| Variable Name | Description |
| --- | --- |
| Count | Count of ROH per individual and population |
| Count Var. | Variance of count |
| Gap | Distance between two adjacent ROH, median per individual and population |
| Gap Var. | Variance of gap |
| Length | Length of ROH, median per population |
| Length Var. | Variance of length |
| $F_{ROH}$ | $F_{ROH}$ (proportion of genome covered by ROH, mean per population) |
| $F_{ROH}$ Var. | Variance of $F_{ROH}$ |
| Selfing Rate | Selfing rate, from 0 (obligate outcrossing) to 1 (obligate selfing) |
| Tajima's $D$ | Tajima's $D$ |
| $F$ | Inbreeding coefficient |
| Abs. Error | Absolute error of prediction for selfing rate |

**Fig. S1:**
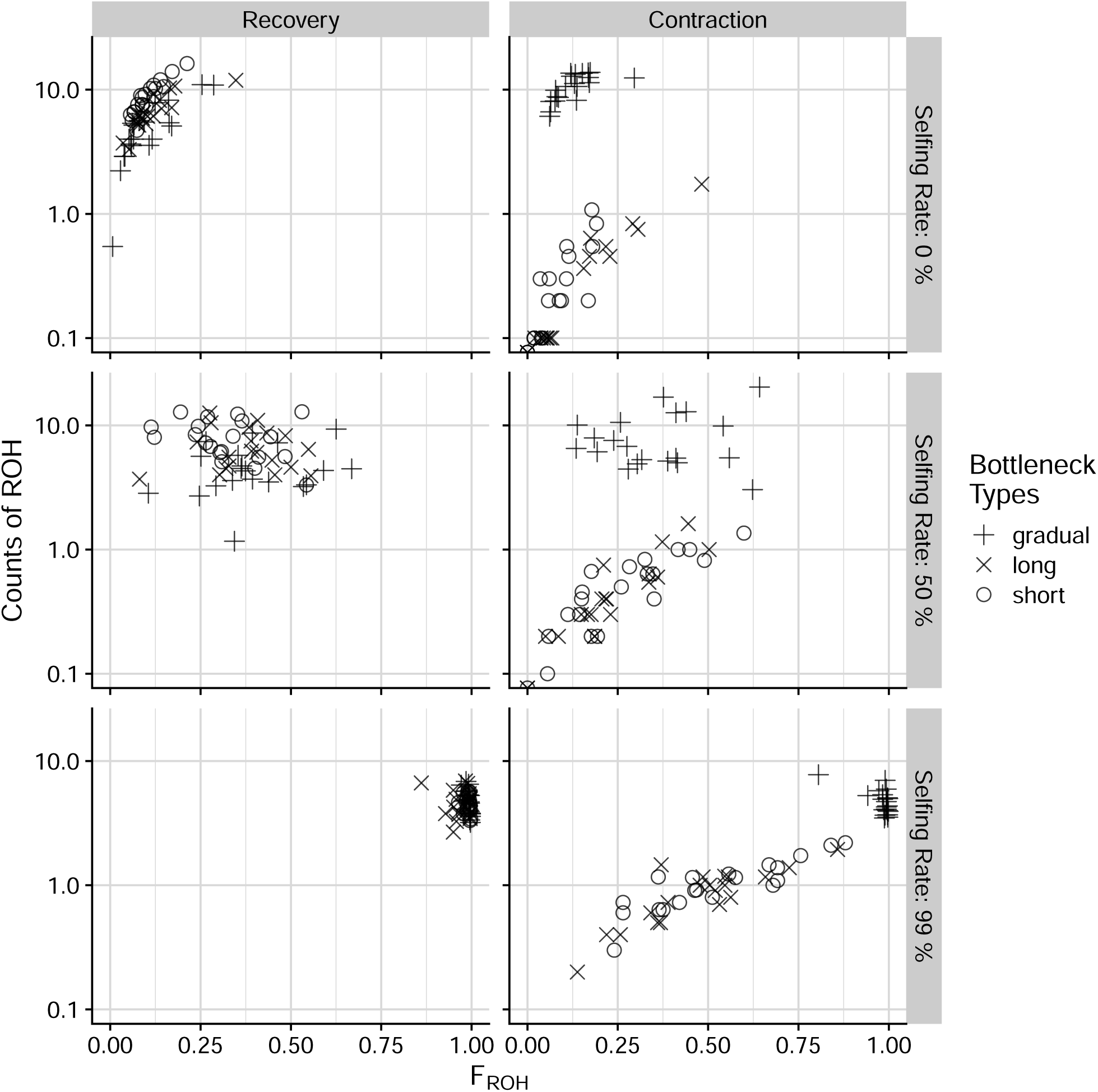
Count of ROH and *F_ROH_*are shown for different bottleneck types and sampling timepoints. For clarity, only samples of an initial population size of *N* = 1000 are shown. ‘Contraction’ populations, sampled during the bottleneck are in the left column, and ‘recovery’ populations, sampled 5000 generations after the contraction started are shown on the right.

**Fig. S2:**
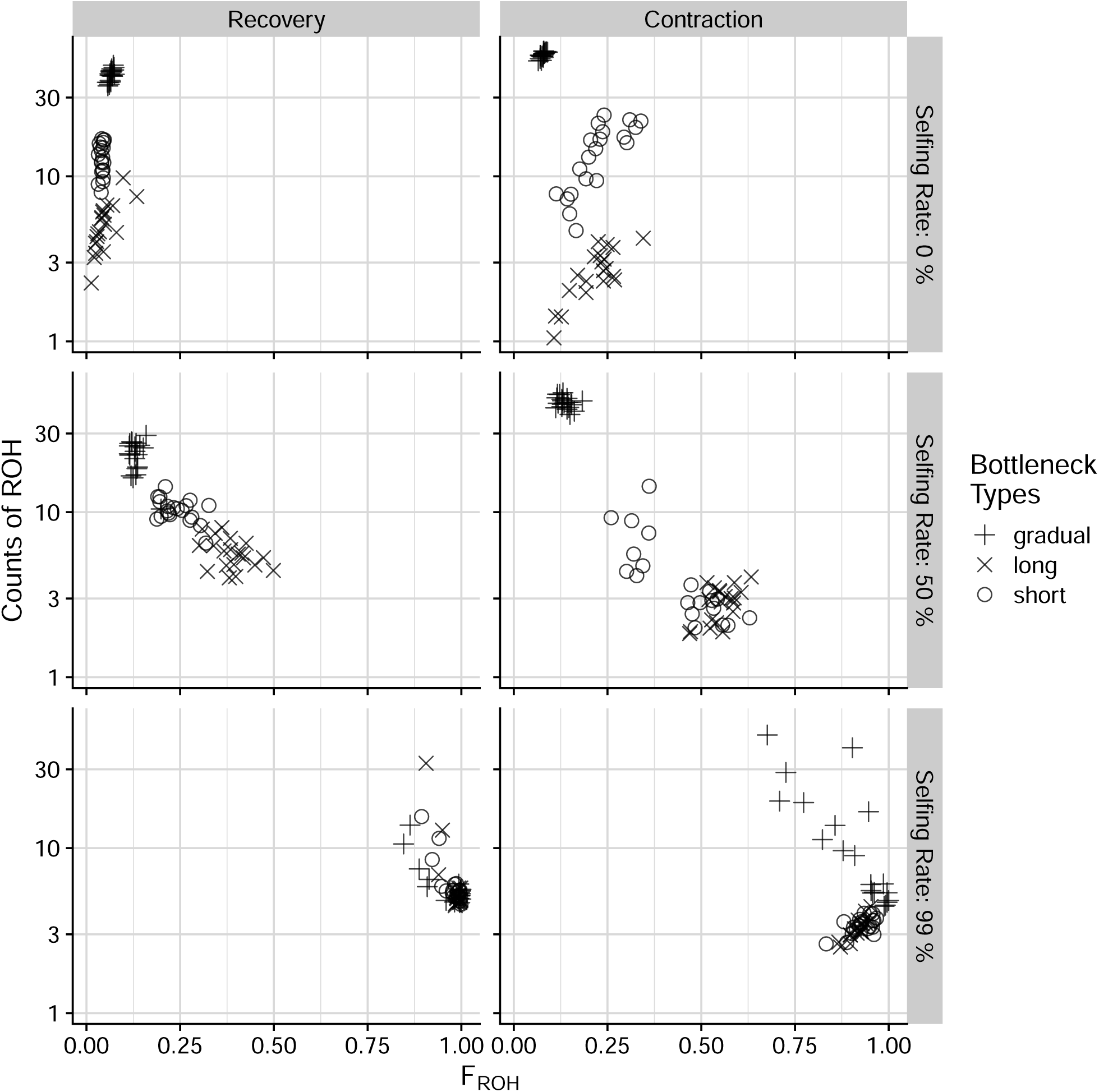
Count of ROH and *F_ROH_* are shown for different bottleneck types and sampling timepoints; similar to Fig. S1, but for an initial population size of *N* = 10000.

**Fig. S3:**
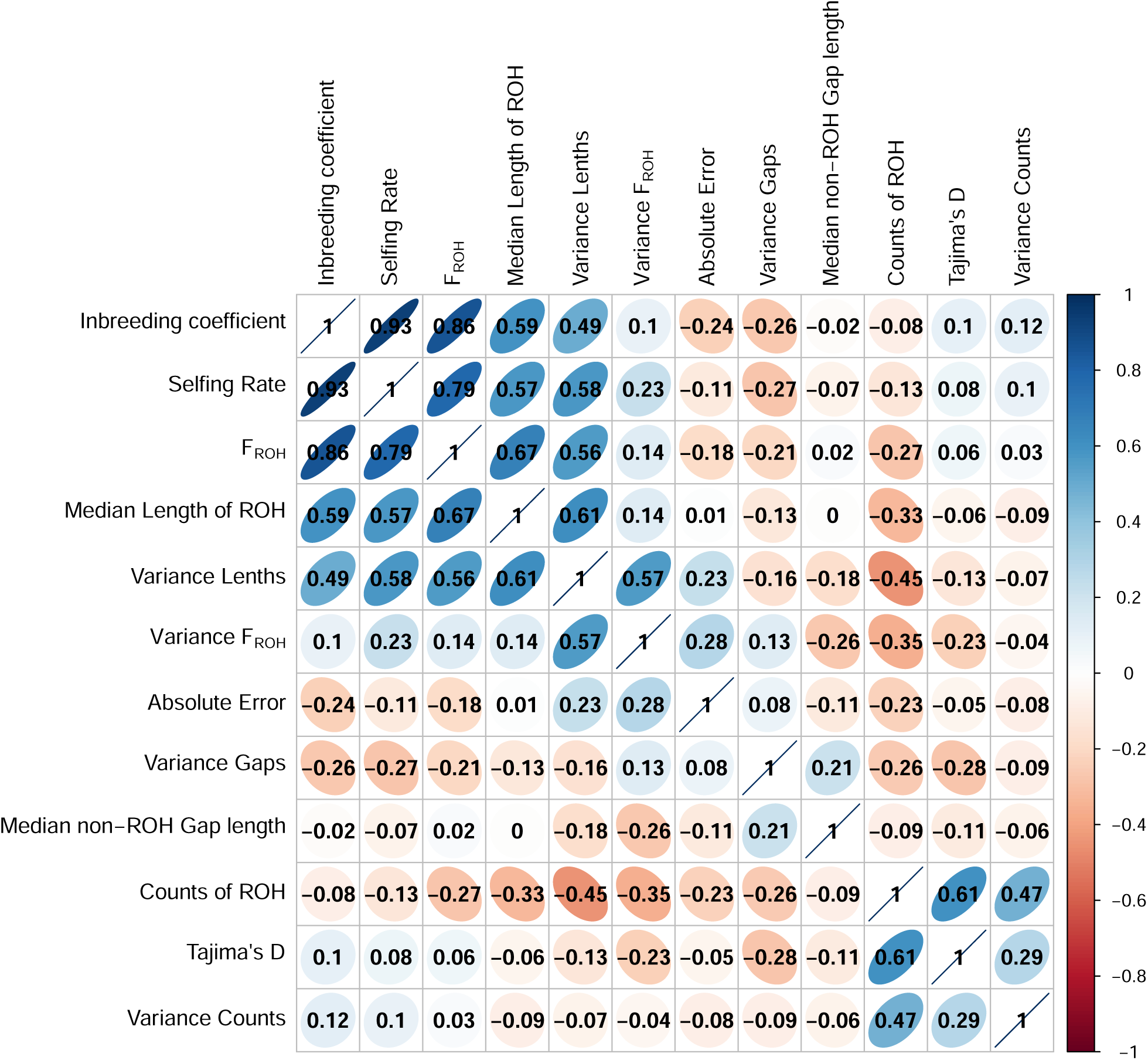
Pairwise comparison of correlation coefficients for predictor variables derived from runs of homozygosity, and inbreeding coefficients (*F*), Tajima’s *D* and the absolute error of the predictions for selfing rate, using the full RF model. The data is from the validation dataset. Variables are ordered according to the angular order of the eigenvectors (Friendly 2002). The figure was created using the R package corrplot (Wei and Simko 2021).

**Fig. S4:**
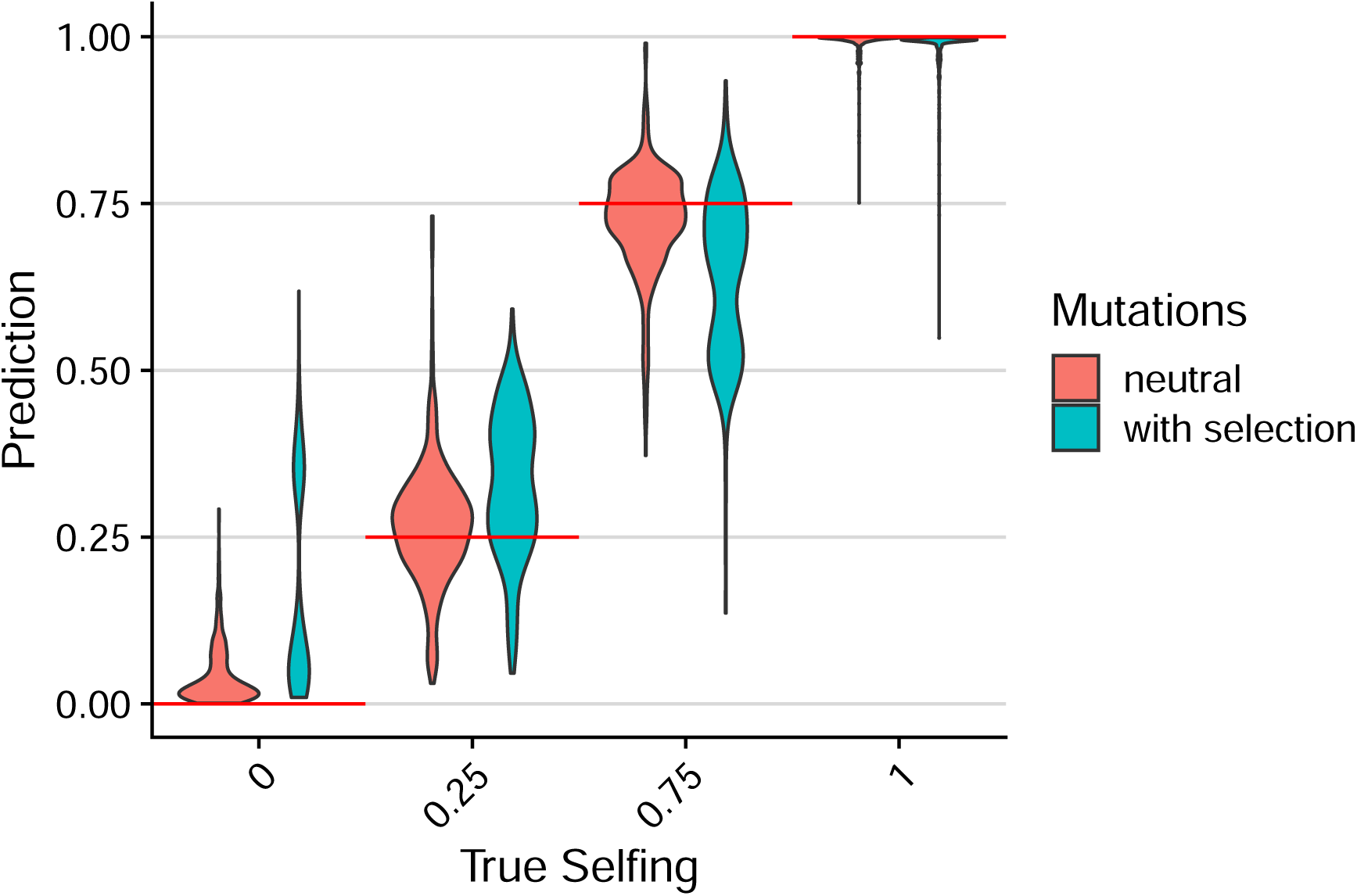
Similar to Fig. 2E, we compare true selfing rates in the test data with predictions of the RF-full model. Compared to simulation scenarios that only included neutral mutations (red), cases with selection (blue) result in predictions biased towards intermediate selfing rates in some scenarios. Red lines are true selfing rates.

**Fig. S5:**
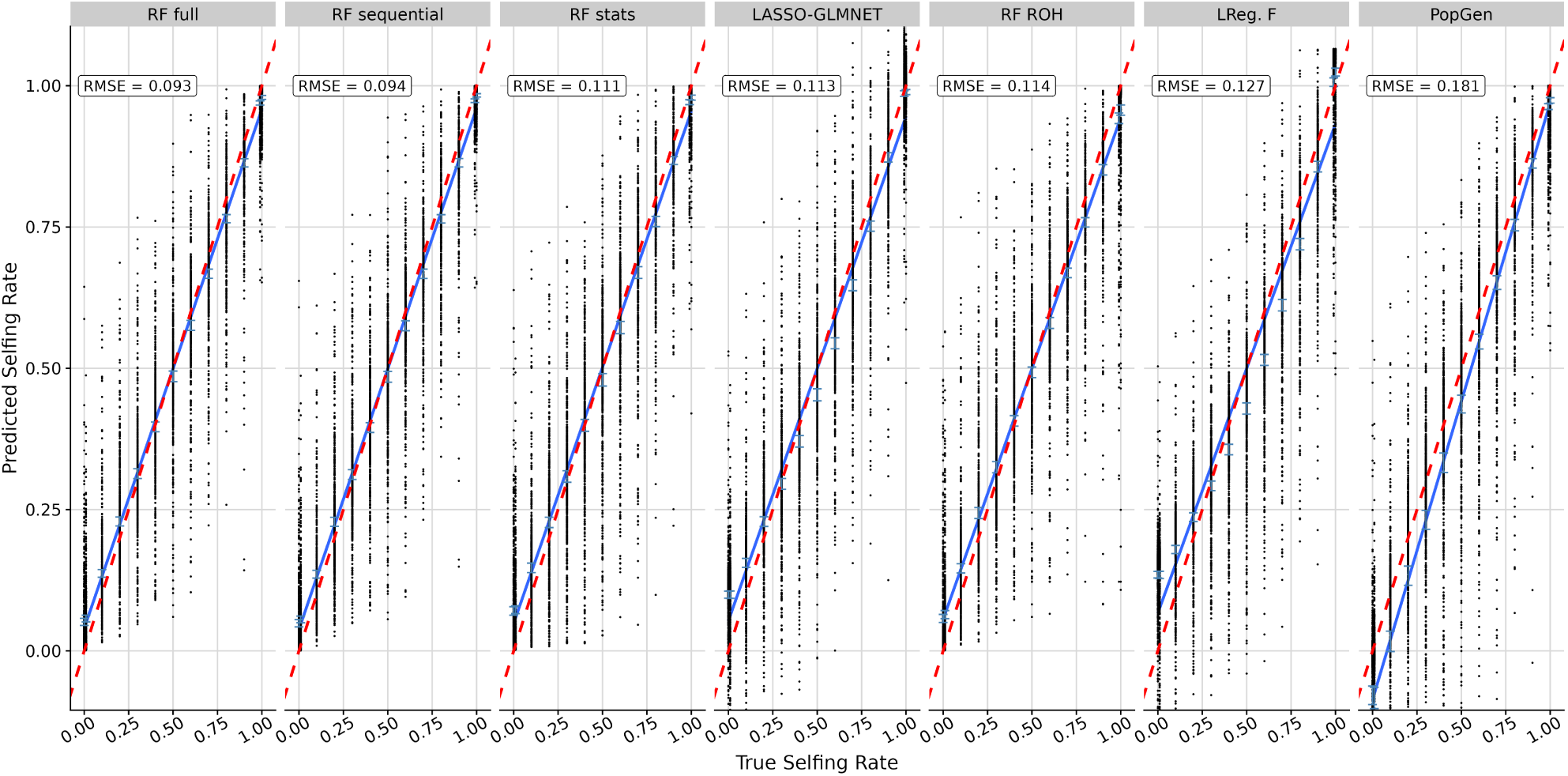
Correlation of true selfing rates and predictions in the validation set for every model. Plot facets are ordered according to RMSE ranked from lowest to highest. Points refer to single population selfing rates (true versus predicted), blue lines are linear fits comparing true and predicted values, and red dashed lines are identity lines, representing perfect predictions.

**Fig. S6:**
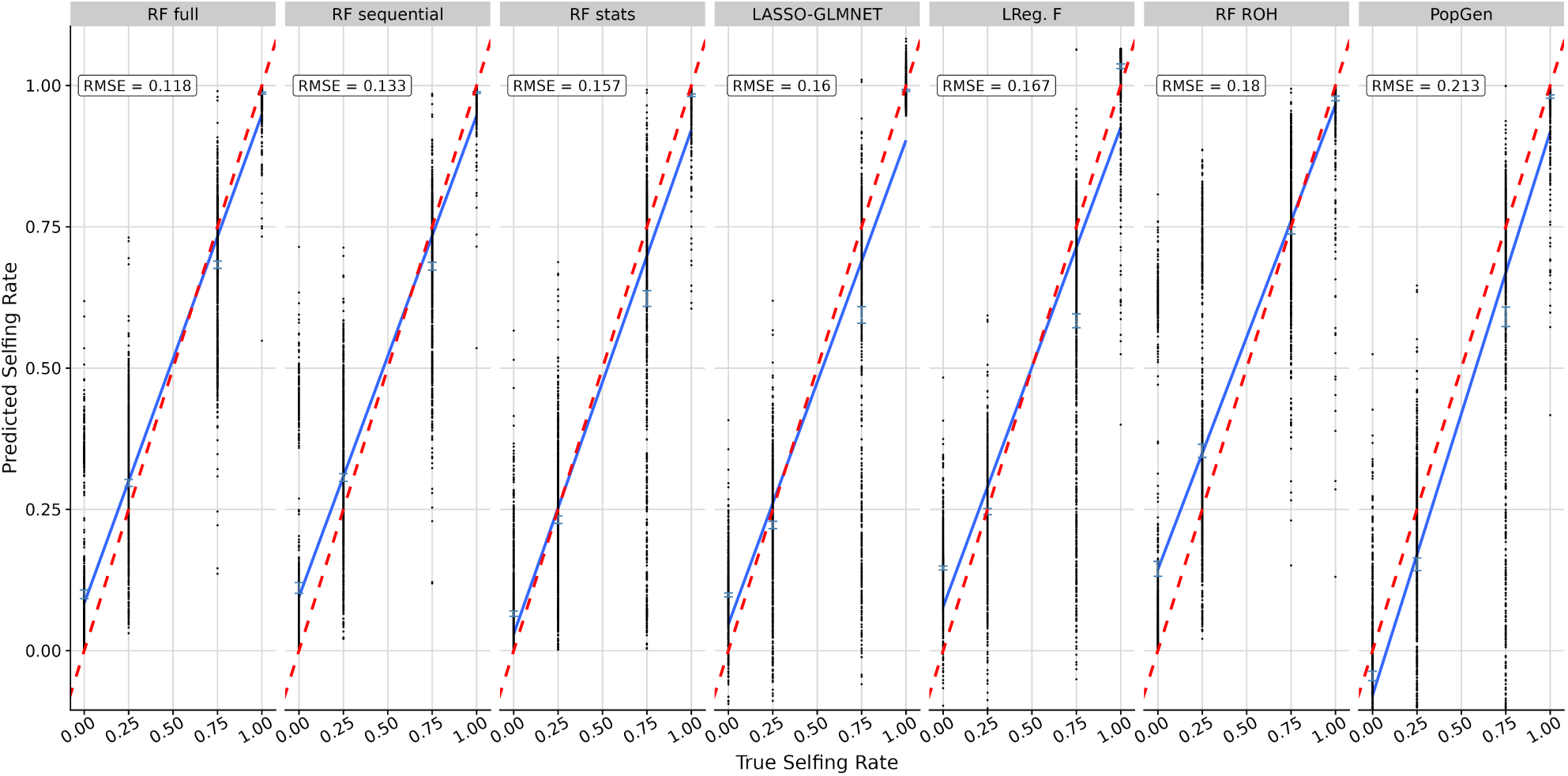
Correlation of true selfing rates and predictions in the unseen test set for every model. Plot facets are ordered according to RMSE, starting with the model with the lowest error on the left. See Fig. S5 for more information.

**Fig. S7:**
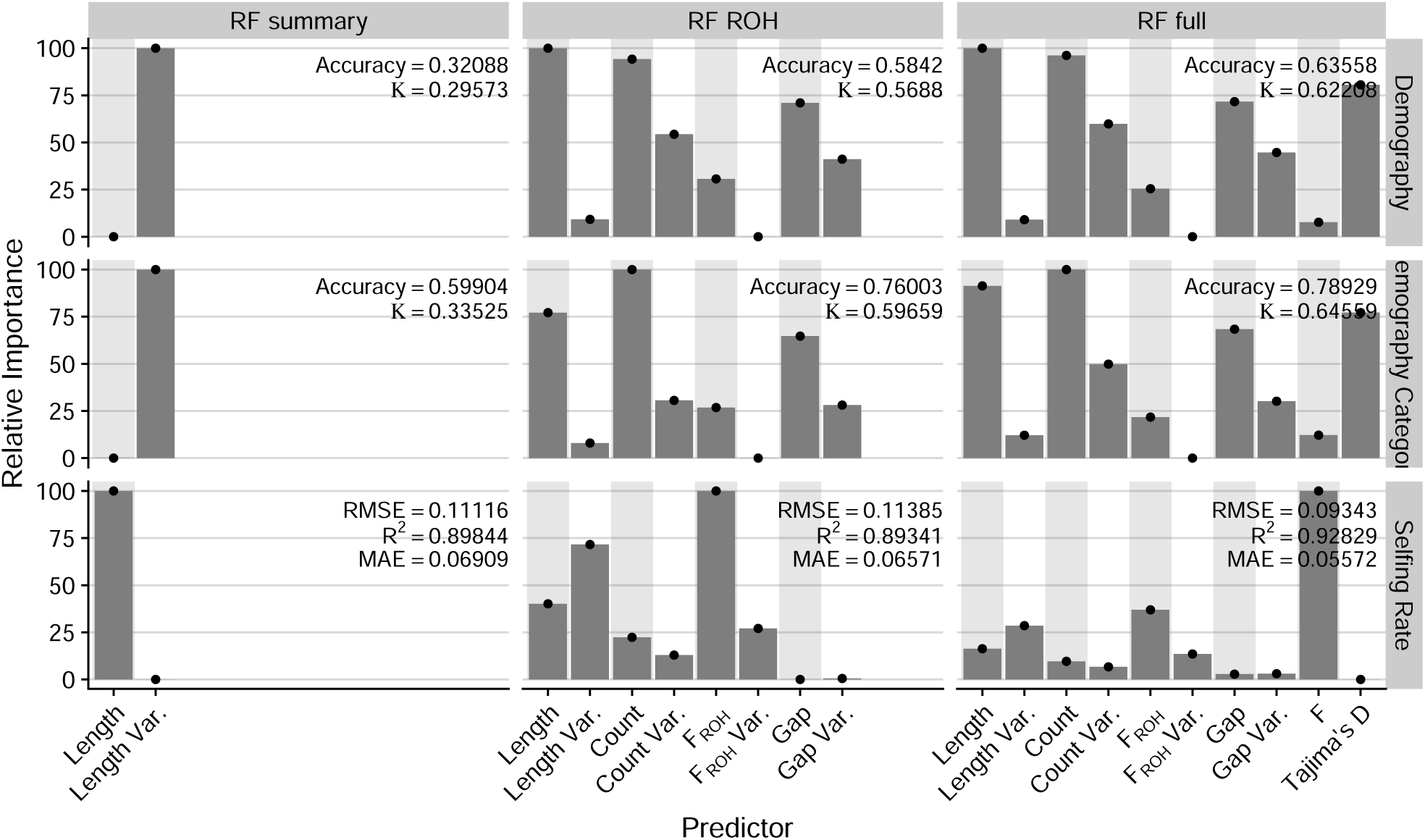
Variable Importance and performance statistics for validation sets for random forest models using summary statistics (left column), ROH statistics (center column) and models with all predictor available, predicting demography (28 categories, top row), a broad category of demography (3 classes, center row) or selfing rate (bottom row).

**Fig. S8:**
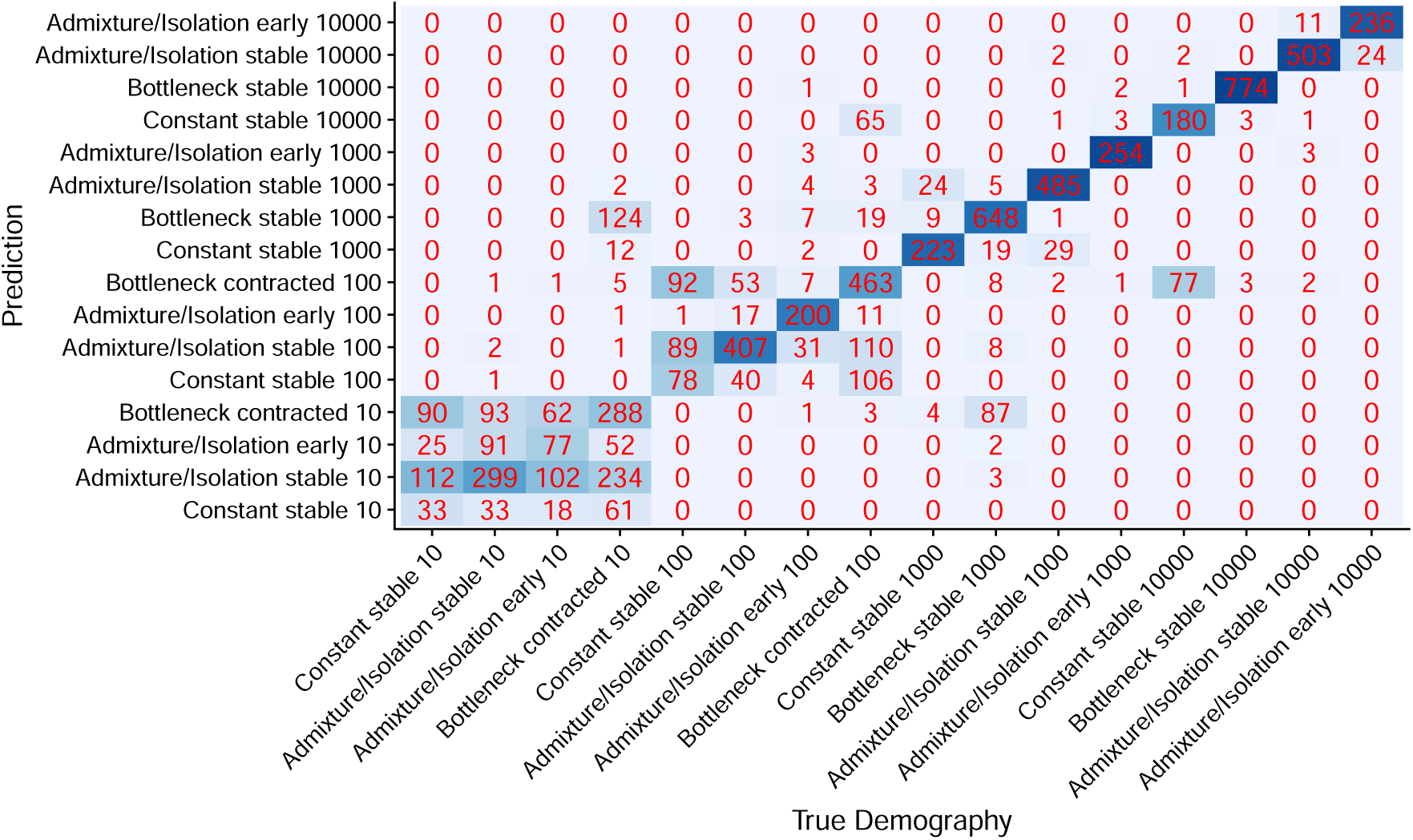
Confusion matrix for the full RF model using the test dataset. Demographic categories are grouped by their type, time point of sampling (early, stable, contracted) and N of this time point. For simplicity, not all 28 demographic scenarios are shown as separate categories in this plot.

**Fig. S9:**
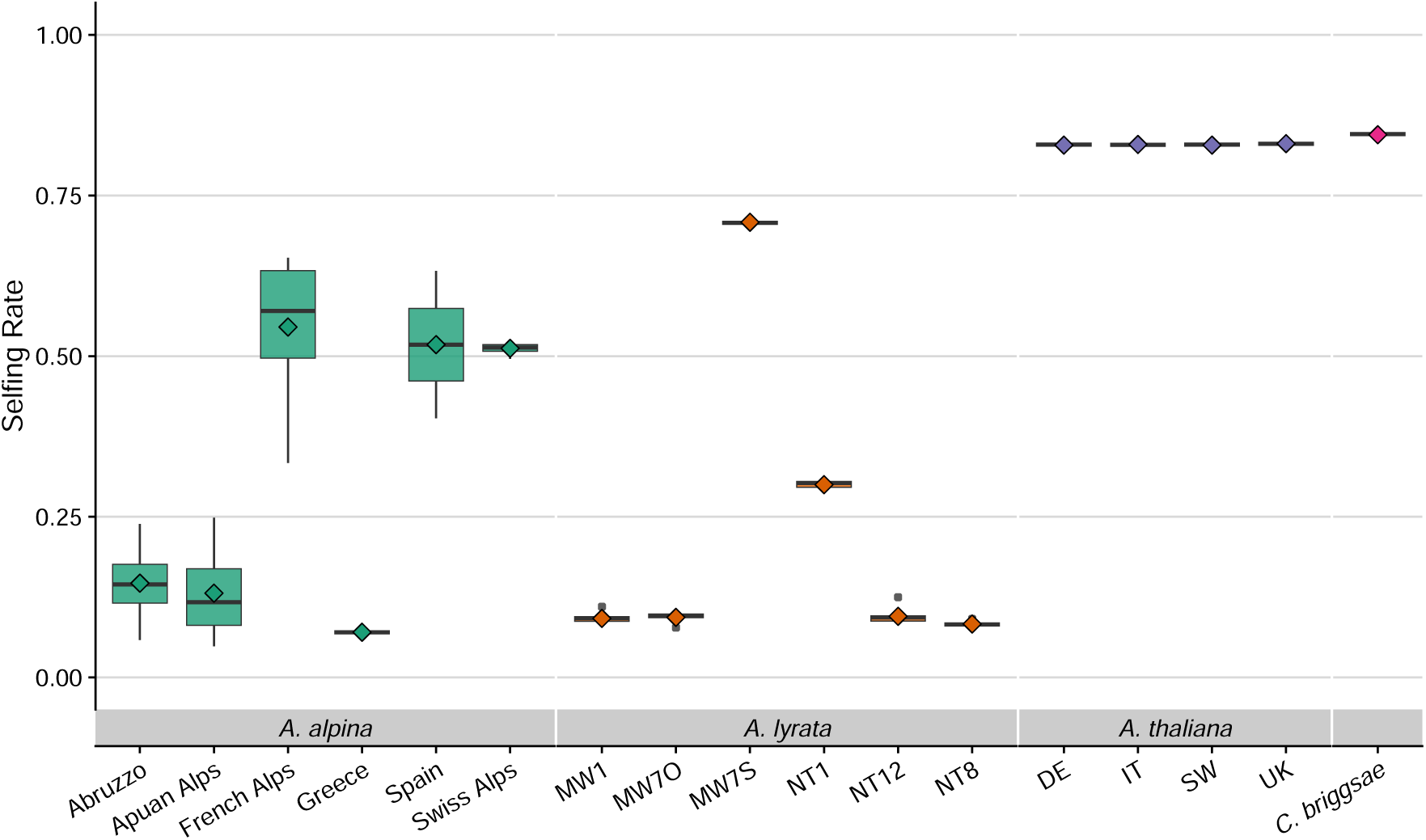
Estimated selfing rates for the full Random Forest model for all four analysed species. Diamonds are group means, estimates for *A. alpina* are per population, and plotted grouped by geographical origin of these populations. Other populations are individual boxes.

**Fig. S10:**
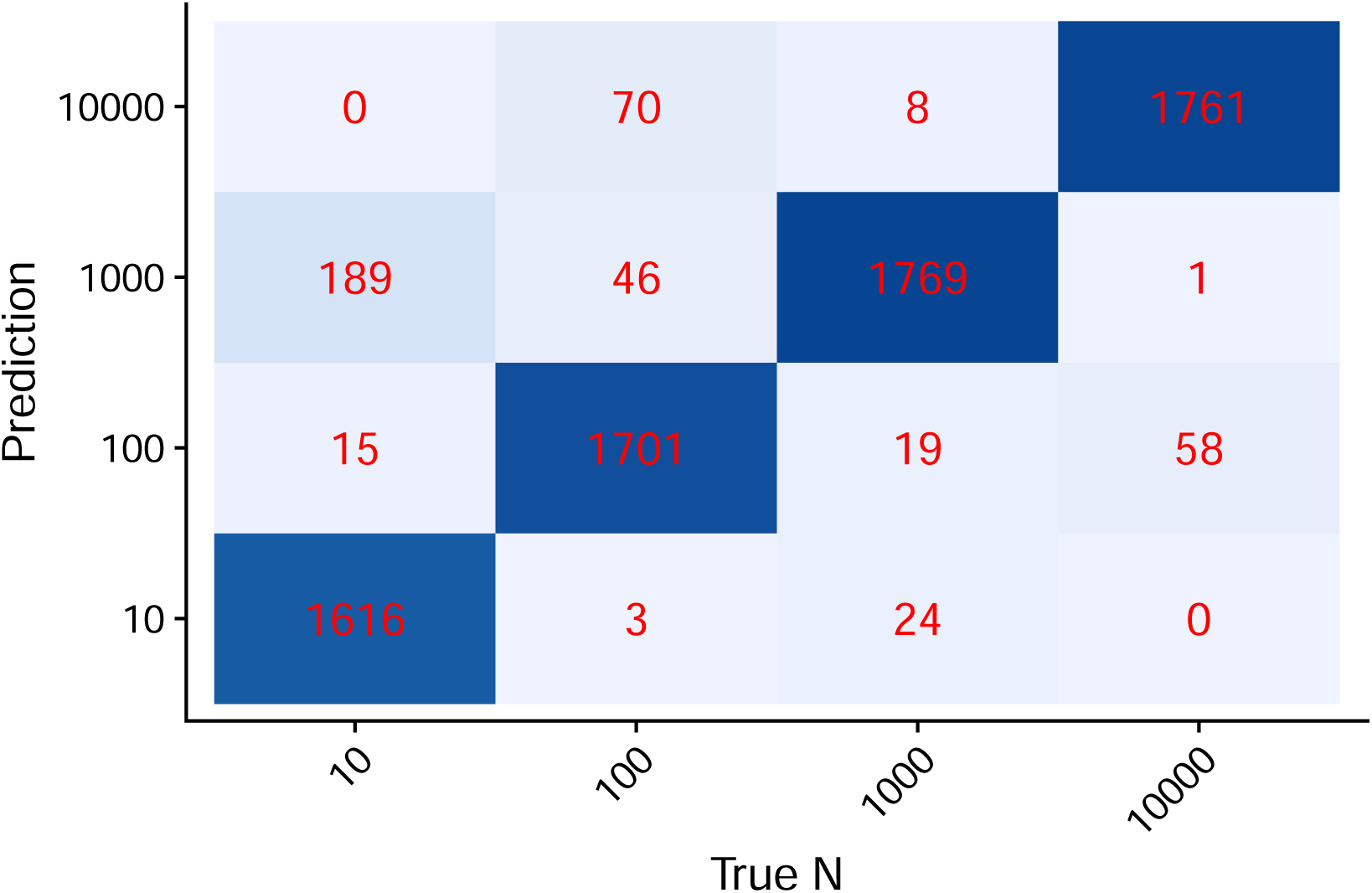
Confusion matrix for the first step in the sequential prediction, the inference of *N*. The accuracy was 0.941 predicting *N* using this model.

**Fig. S11:**
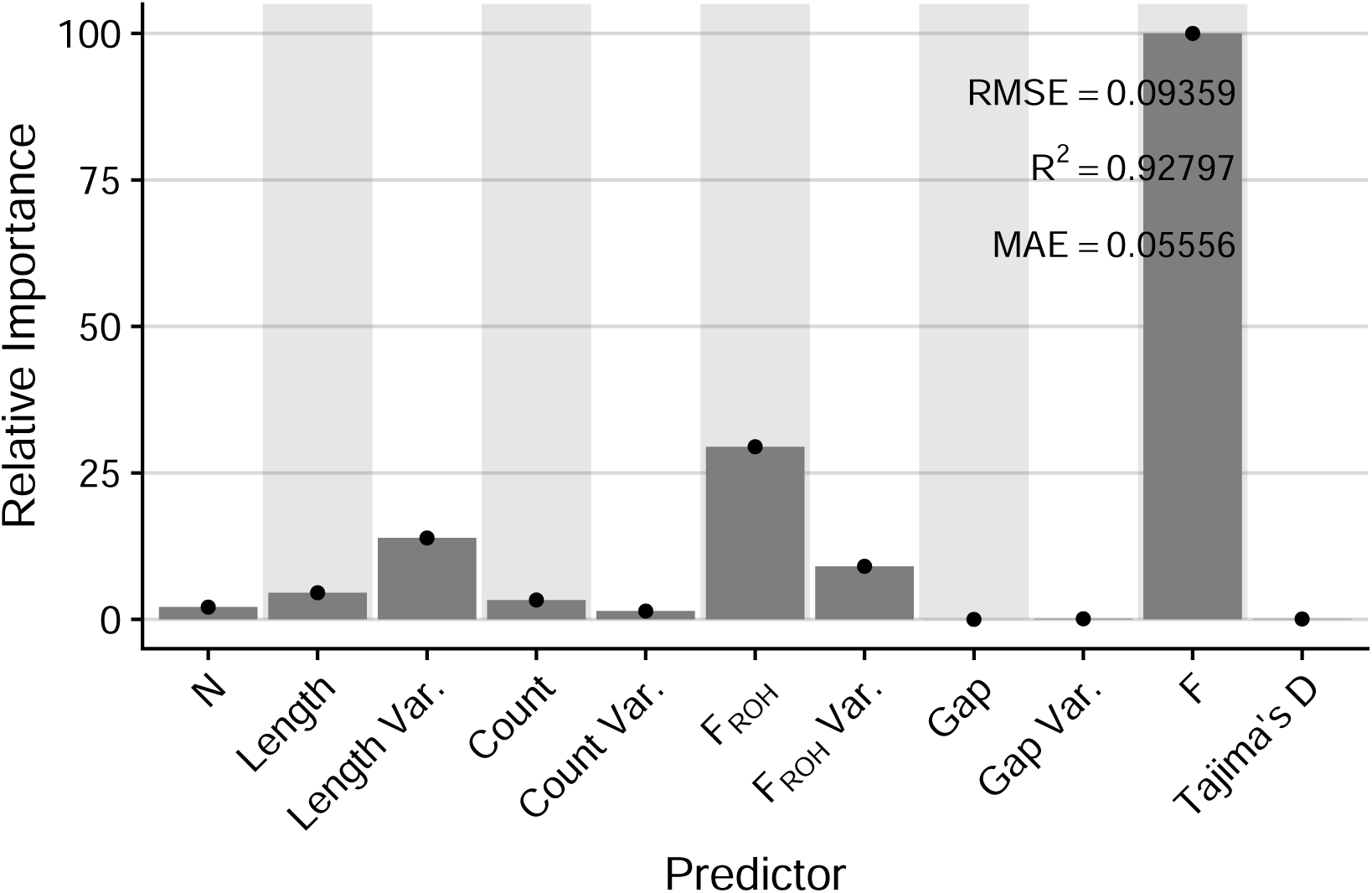
Variable importance and performance metrics of the sequential random forest model to predict selfing rate for the validation data.

